# An expanded 2-pyridinecarboxaldehyde (2PCA)-based chemoproteomics toolbox for probing protease specificity

**DOI:** 10.1101/2023.02.12.528234

**Authors:** Haley N. Bridge, Clara L. Frazier, Amy M. Weeks

**Affiliations:** Department of Biochemistry, University of Wisconsin – Madison, Madison, WI, USA 53706

## Abstract

Proteomic profiling of protease-generated N termini, or N terminomics, provides key insights into protease function and specificity. However, current N terminomics technologies have sequence limitations or require specialized synthetic reagents for N-terminal peptide isolation. Here, we introduce an expanded N terminomics toolbox that is based on 2-pyridinecarboxaldehyde (2PCA) reagents. These tools enable efficient enrichment of protein N termini by combining selective N-terminal biotinylation using 2PCA reagents with chemically cleavable linkers for N-terminal peptide recovery. By incorporating a commercially available alkyne-modified 2PCA in combination with Cu(I)-catalyzed azide-alkyne cycloaddition (CuAAC), our strategy eliminates the need for chemical synthesis of N-terminal probes. Using these reagents, we developed PICS2 (<u>P</u>roteomic Identification of <u>C</u>leavage <u>S</u>ites with <u>2</u>PCA reagents) to profile the specificity of subtilisin/kexin-type proprotein convertases (PCSKs). We also implemented CHOPPER (<u>Ch</u>emical enrichment <u>O</u>f <u>P</u>rotease substrates with <u>P</u>urchasable, <u>E</u>lutable <u>R</u>eagents) for global sequencing of apoptotic proteolytic cleavage sites. Based on their broad applicability and ease of implementation, PICS2 and CHOPPER are useful tools that will advance our understanding of protease biology.

## Introduction

Proteolytic cleavage regulates the activity, localization, and lifetime of nearly all human proteins. Although proteases are encoded by ∼2% of human genes^1^ and are an important class of drug targets^2^, the substrate specificity of many proteases remains unknown or incompletely understood. Proteolytic cleavage of a protein results in formation of new N and C termini. Chemoproteomics methods that combine selective isolation of protein N termini with tandem mass spectrometry (LC-MS/MS) are powerful tools for defining exact sites of proteolytic cleavage, enabling protease sequence specificity profiling^3^ and identification of cellular protease substrates^4–8^. However, current N terminomics methods are limited by sequence specificity, low efficiency, and challenges to their implementation that include the need to synthesize specialized reagents.

A necessary feature of enrichment-based N terminomics workflows is selective biotinylation of the N terminus over other primary amines including Lys side chains^3,4,7^. Selectivity is typically achieved using one of two approaches. In the first approach, all primary amines, including both N termini and Lys side chains, are blocked prior to protease digestion to enable biotinylation of proteolytic neo-N termini with an amine-reactive reagent^3^. This approach precludes the analysis of a large group of physiologically important proteases, including trypsin and subtilisin/kexin-type proprotein convertases (PCSKs), that recognize Lys as part of their consensus cleavage sequences^9,10^. An alternative approach for N-terminal biotinylation employs an N-terminally selective peptide ligase enzyme^4,8,11^. This method does not require modification of Lys side chains, but the N-terminal sequences that are biotinylated may be biased based on the ligase’s intrinsic sequence specificity^11^.

Because of their high N-terminal specificity and their low sequence specificity, probes based on 2-pyridinecarboxaldehyde (2PCA) (**1**) are attractive as N terminomics tools to overcome these challenges^7,12^. The specificity of 2PCA reagents for the N terminus over Lys is enforced by the chemical mechanism of modification, which involves initial formation of an N-terminal imine followed by nucleophilic attack of the neighboring amide in the peptide backbone to form a stable cyclic imidazolidinone (**Fig. 1A**)^12^. Because Lys side chains lack a neighboring amide group positioned for cyclization, they do not form stable adducts with 2PCA. In principle, 2PCA modification can occur on any N-terminal sequence except those with Pro in the second position, which lack the backbone amide NH needed for cyclization. Biotin-2PCA has been previously applied to enrich substrates of dipeptidyl peptidases in a workflow termed CHOPS (<u>Ch</u>emical enrichment <u>O</u>f <u>P</u>rotease <u>S</u>ites), providing a foundation for unbiased identification of protease substrates in complex samples^7^. However, adoption of 2PCA reagents as protease substrate probes is limited by the need to synthesize the required biotinylated probe and by the absence of a cleavable linker for efficient elution of N-terminal peptides prior to LC-MS/MS. Additionally, the sequence specificity of 2PCA modification in complex samples has not been fully characterized.

**Figure 1.**
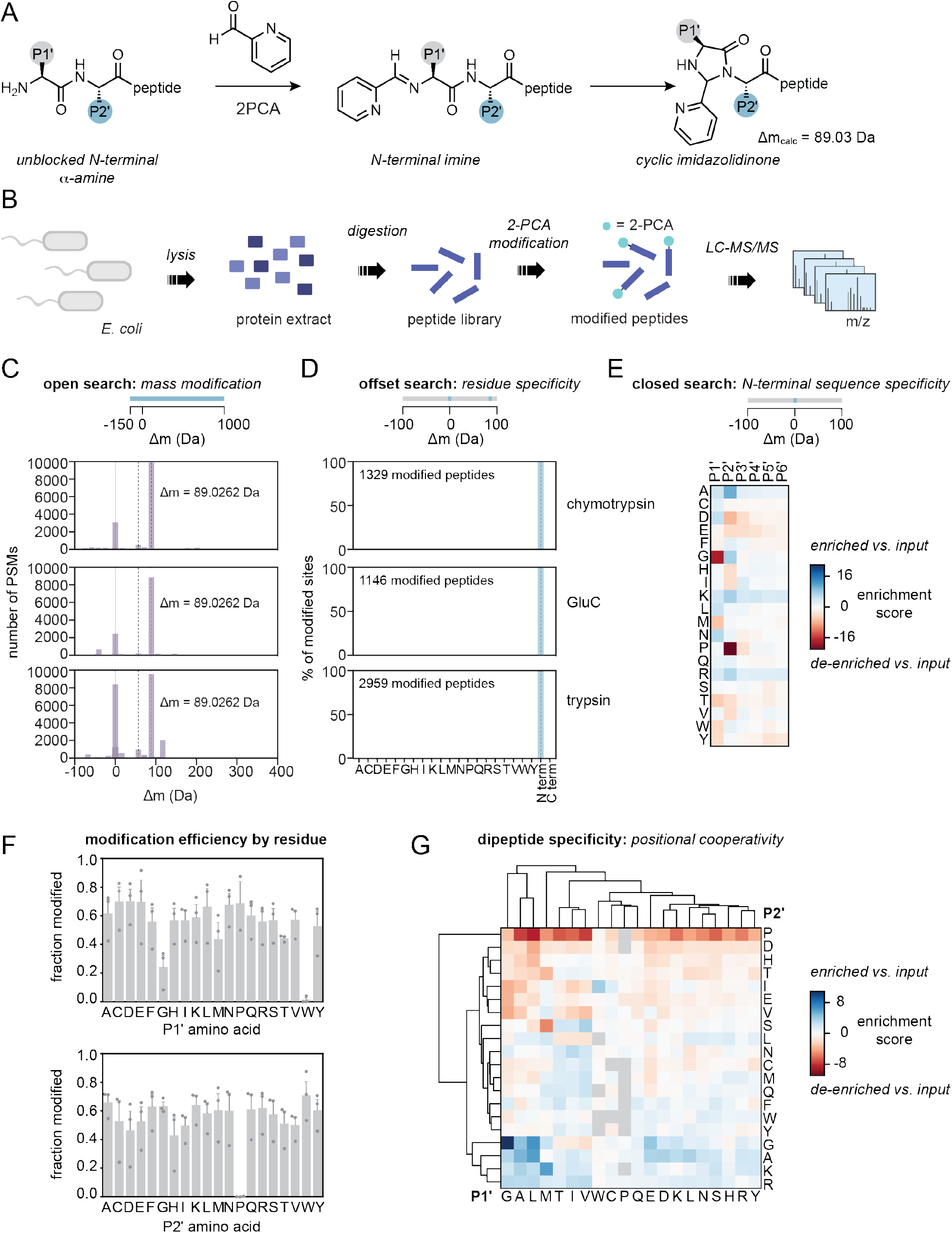
Deep profiling of 2PCA specificity using proteome-derived peptide libraries. (A) 2-pyridinecarboxaldehyde selectively modifies the N terminus of peptides, initially forming an N-terminal imine and then cyclizing to for a cyclic imidazolidinone. (B) Workflow for 2PCA specificity profiling. Proteome-derived peptide libraries were generated with *E. coli* protein extract by protease digestion, treated with 2PCA, and analyzed using LC-MS/MS to determine sites of modification. Sequence logos for input libraries are shown in Fig. S1. (C) Open database search to determine the number of 2PCA modifications that occur per peptide. (D) Offset database search to determine the residue specificity of 2PCA modification. (E) Closed database search to determine the sequence specificity of 2PCA modification. Results for individual libraries are shown in Fig. S2 and MS/MS spectra for putative modified peptides with P2‘ Pro are shown in Fig. S3. (F) Modification of efficiency of 2PCA by the residue in the P1‘ position (top) or the P2‘ position (bottom). (G) Analysis of P1‘-P2‘ pairwise residue interactions on the efficiency of 2PCA modification efficiency. Heatmaps for other pairwise interactions are shown in Fig. S4.

Here, we report an expanded toolbox of N terminomics probes based on 2PCA reagents. We fully defined the sequence specificity and biases of 2PCA modification and optimized modification conditions using proteome-derived peptide libraries. We designed a biotin-2PCA with a cleavable linker to enhance the efficiency of peptide elution and incorporated a commercially available alkyne-modified 2PCA in combination with Cu(I)-catalyzed azide-alkyne cycloaddition (CuAAC) to enable implementation of 2PCA-based N terminomics with off-the-shelf reagents. We applied these probes to develop a positive-enrichment strategy to define protease sequence specificity in proteome-derived peptide libraries that we term <u>P</u>roteomic Identification of <u>C</u>leavage <u>S</u>ites with <u>2</u>PCA reagents (PICS2). We deployed PICS2 to profile the sequence specificity of the subtilisin/kexin-type proprotein convertases (PCSKs) Kex2, furin, and PCSK2, biologically important proteases whose sequence specificity cannot be accurately characterized with existing methods. We also introduce <u>Ch</u>emical enrichment <u>O</u>f <u>P</u>rotease substrates with <u>P</u>urchasable, <u>E</u>lutable <u>R</u>eagents (CHOPPER), a method for enrichment of protease substrates from cell lysates that uses commercially available, selectively elutable probes. We applied CHOPPER to profile proteolytic cleavage events in etoposide-induced apoptosis, leading to the identification of 112 previously unknown putative caspase cleavage sites in 95 proteins. Because they are easy to implement and broadly applicable, PICS2 and CHOPPER greatly expand the toolbox for proteome-wide study of protease cleavage sites to advance our understanding of proteolytic signaling pathways.

## Results

### Proteome-derived peptide libraries for deep profiling of 2PCA specificity

N-terminal enrichment strategies enable isolation of the prime side peptide product (P’, C-terminal to the scissile bond) that results from protease cleavage and enable inference of identify of the cleaved protein and the sequence of the nonprime side peptide product (P; N-terminal to the scissile bond)^4,7,13^. Although an ideal N-terminal modification reagent for N terminomics applications would react with all N-terminal sequences in an unbiased manner, all selective N-terminal modification methods reported to date have sequences biases^14^. Understanding the sequences biases intrinsic to each N-terminal probe is therefore important for determining its suitability for any particular application. Previous characterization of 2PCA specificity focused on varying the N-terminal amino acid of the sequence XADSWAG, where X is a variable amino acid^12^. These studies found that most peptides were modified with similar efficiencies except for those with X = G or X = P, which were modified to a lower extent. Because it focused on a single peptide sequence, this study did not address the effect of positions beyond the N-terminal amino acid on 2PCA modification efficiency, including the second amino acid, whose backbone amide is proposed to participate in the reaction. We therefore set out to more comprehensively characterize 2PCA specificity.

To characterize 2PCA specificity in depth, we used a mass spectrometry-based assay inspired by proteomic identification of ligation sites (PILS), which was previously developed for comprehensive and quantitative characterization of peptide ligase N-terminal specificity (**Fig. 1B**)^11^. We generated three N-terminally diverse proteome-derived peptide libraries by digesting *E. coli* protein extracts with three different proteases with distinct P1 specificities: trypsin (P1 = K or R), GluC (P1 = E or D), and chymotrypsin (P1 = F, L, W, or Y). In combination, these libraries represent every possible amino acid in the first six N-terminal positions (P1‘-P6‘) and nearly all 400 possible amino acids combinations at P1‘-P2‘ (**Fig. S1**). We incubated each peptide library with 10 mM 2PCA at pH 7.5 for 4 h at 37ºC and then analyzed the 2PCA-treated peptide libraries by LC-MS/MS to determine (1) the number of 2PCA modifications on each modified peptide; (2) the site specificity of the 2PCA modification; (3) the N-terminal sequence specificity of the 2PCA modification; (4) the positional cooperativity of the 2PCA modification; and (5) the efficiency of modification of each N-terminal residue.

The initial report of 2PCA reagents for N-terminal modification suggested that some N termini might harbor more than one 2PCA modification^12^. To empirically determine the mass modification(s) attributable to 2PCA treatment, we used the open search workflow of the FragPipe proteomics pipeline^15,16^ for unbiased analysis of 2PCA modifications across the 2PCA-treated peptide libraries (**Fig. 1C**). The open search workflow allows a wide precursor mass tolerance (-150 Da to 500 Da), enabling identification of modifications based on the empirical data without the need to specify them during database search. Using this method, we found that Δm = 89.0262 Da was the most abundant mass modification in all three peptide libraries. This finding supports the hypothesis that a single 2PCA modification (Δm_calc_ = 89.0265 Da) occurs on each modified peptide. No mass modification corresponding to double 2PCA modification was observed.

Although 2PCA possesses an aldehyde that may react reversibly with both N-terminal α amines and lysine side chains, stable 2PCA modification is proposed to be restricted to N termini based on the need for an adjacent amide bond to participate in cyclic imidazolidinone formation (**Fig. 1A**). In support of this proposal, previous work demonstrated that peptides with a blocked N terminus, such as hypertrehalosemic neuropeptide, which has a pyroglutamate residue at the N terminus, cannot be modified with 2PCA^12^. Additionally, 2PCA modified a panel of protein substrates only once per polypeptide chain^12^. To test this proposal in the context of a wide variety of N-terminal peptide sequences, we globally analyzed 2PCA selectivity for all amino acid side chains and N and C termini using the offset search strategy of the FragPipe proteomics pipeline (**Fig. 1D**)^15,16^. This approach allows the user to specify a mass modification without the need to restrict the residues on which the modification may occur. Offset search of the three 2PCA-modified peptide libraries with a mass modification of 89.0265 Da revealed that 2PCA exhibits absolute chemoselectivity for peptide N termini, with no modifications observed on lysine or any other side chain.

We next evaluated whether 2PCA exhibits N-terminal sequence specificity (**Fig. 1E, Fig. S2**). Based on the mechanism of cyclic imidazolidinone formation, 2PCA is unable to modify peptides with Pro in the second position as they lack the amide NH group required for cyclization12. Beyond this mechanism-based specificity, the efficiency of 2PCA modification of different peptide sequences has not been investigated in depth. To enable identification of the maximum number of 2PCA-modified peptides, we use a closed search strategy in which the search for the 2PCA modification was restricted to the N terminus. In total, we identified 28,167 peptides across the tryptic, GluC, and chymotryptic peptide libraries, of which 14,735 (52%) were 2PCA-modified. To examine 2PCA specificity in greater detail, we compared the position-specific frequencies of each amino acid in the 2PCA-modified peptides and in the input peptide libraries (**Fig. 1E**). As predicted by the proposed mechanism for 2PCA modification, we observed a strong bias against 2PCA modification of peptides with Pro in the second position. Of 900 peptides with Pro in position 2 identified by database searching across the three libraries, only two (0.2%) were identified as 2PCA-modified (**Fig. 1F**). Each of the putative 2PCA modified peptides with Pro in position 2 was identified with one peptide spectrum match, and both had relatively low cross-correlation (XCorr) scores (0.88 and 1.06, respectively) and either zero or one fragment ion matches that are specific to the modified peptide (**Fig. S3**). Consistent with previous studies^12^, we also observed a bias against 2PCA modification of peptides with glycine in the first position (**Fig. 1E**), with 356 of 1,669 (21%) of peptides with glycine at position 1 bearing a 2PCA modification compared to 52% of peptides in the population (**Fig. 1F**). However, in contrast to previous work that showed that X = P inhibited 2PCA modification of the peptide XADSWAG, we observed no bias against modification of peptides with Pro in position 1. Beyond the biases against modification of Gly in position 1 and Pro in position 2, we did not observe substantial preference for or against 2PCA modification of any other amino acid in the first two positions, and no strong sequence specificity was observed beyond the first two positions of the peptide.

In enzymes including proteases and peptide ligases, substrate amino acid subsites often exhibit subsite cooperativity, with the identity of the amino acid in one position of the substrate influencing the enzyme’s ability to act on substrates containing another amino acid in a different position^11,17^. However, the potential for such interactions in the context of 2PCA has not been considered. We therefore examined how the efficiency of 2PCA modification for a peptide with a given amino acid at one position is influenced by the identity of the amino acid at other positions within the peptide (**Fig. 1G, Fig. S4**). We analyzed enrichment or de-enrichment of each pairwise sequence combination at positions 1 through 6 of modified peptides compared to the input peptide libraries. We found that the identity of the amino acid at position 2 has a strong influence on whether peptides with Gly at position 1 are efficiently modified (**Fig. 1G**). Although the total population of peptides with N-terminal Gly is modified with low efficiency (21%), for the subset of peptides that also have Gly in the second position, the efficiency of labeling is much higher. In total, 72 out of 111 peptides (64%) beginning with Gly-Gly were 2PCA modified, making Gly-Gly the most efficiently labeled N-terminal dipeptide sequence. Similar but smaller effects were observed for Gly-Ala peptides (58 of 185 peptides modified, 31.4%), Gly-Lys peptides (27 of 85 peptides modified, 31.8%) and Gly-Arg peptides (27 of 74 peptides modified, 36.5%). We also observed, as predicted based on the mechanism of 2PCA modification, that modification of peptides with any amino acid at position 1 is suppressed by the presence of proline at position 2. Consistent with the lack of sequence specificity observed beyond position 2 in position-specific analysis of 2PCA-modified peptides, we did not observe any pairwise effects beyond the second amino acid (**Fig. S4**).

### Optimizing 2PCA modification of proteomic samples

Encouraged by the broad sequence compatibility and N-terminal selectivity revealed by deep profiling of 2PCA specificity, we set out to optimize the efficiency of 2PCA N-terminal modification for proteomics applications (**Fig. 2A**). We first examined the impact of time, temperature, and buffer concentration on 2PCA modification of an N-terminally diverse proteome-derived peptide library (**Fig. 2BC**). We performed the 2PCA modification reaction for either 4 h or 20 h over a range of temperatures between 37ºC and 75ºC with either 10 mM or 50 mM sodium phosphate, pH 7.5. No substantial effects of buffer concentration were observed. For the 4 h reactions, no substantial difference in labeling efficiency was observed over the range of temperatures (**Fig. 2B**). However, for the 20 h reaction time, the fraction of peptides modified began to decrease at 65ºC and 75ºC, a change which may be attributable to reversal of the 2PCA modification (**Fig. 2C**). Our results demonstrate that 2PCA labeling for 4 h at physiological temperature (37ºC) produces a similar extent of modification to labeling at higher temperatures for longer times. Additionally, these experiments show that prolonged reaction times may be disadvantageous for maximizing the extent of 2PCA modification.

**Figure 2.**
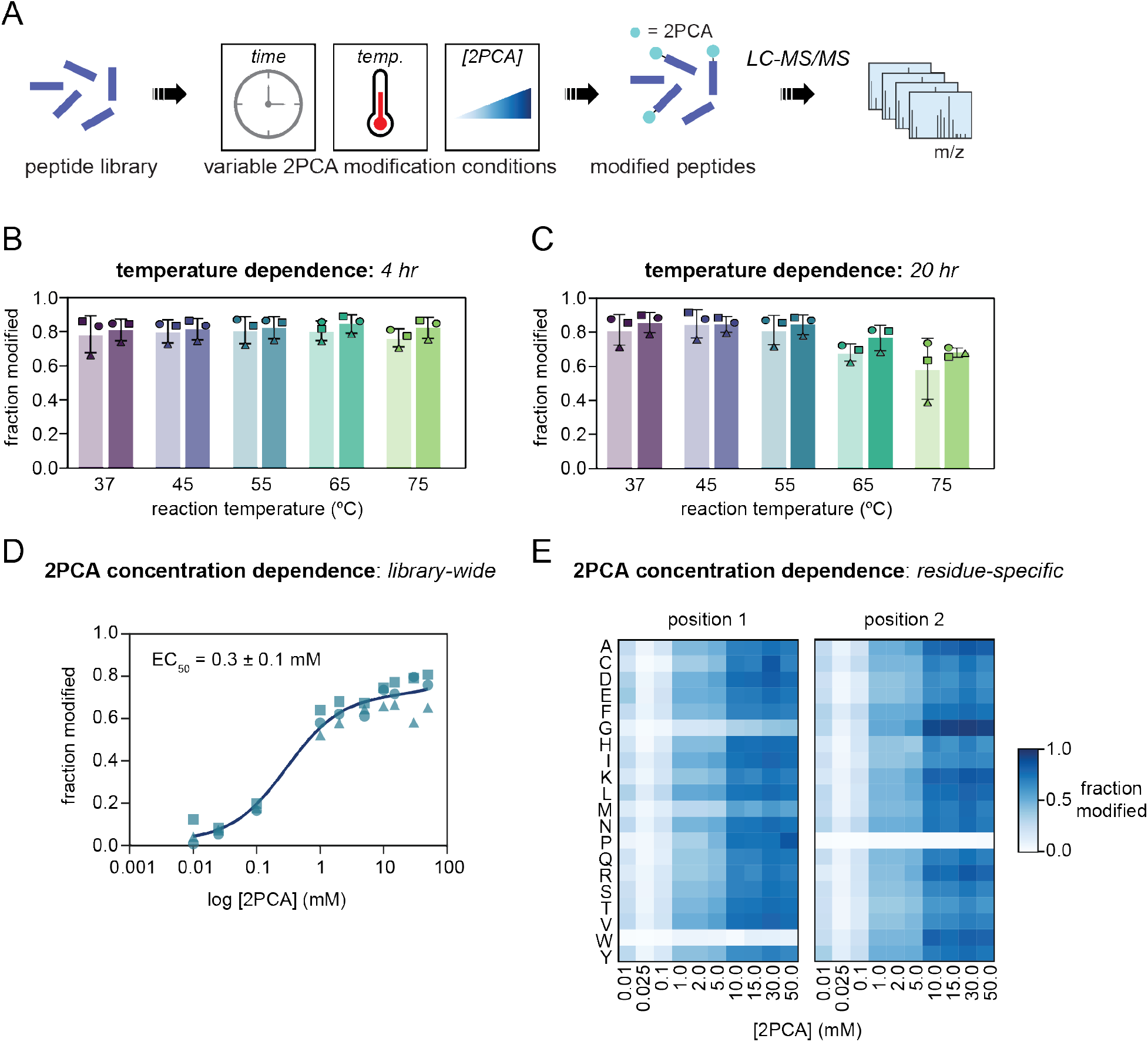
2PCA Labeling Reaction Optimization. (A) Workflow for labeling *E. coli* peptide libraries with 2PCA with variable reaction time, temperature, or 2PCA concentration. (B) Fraction of peptide libraries labeled with 2PCA at 4 h over a range of temperatures from 37ºC to 75ºC. Samples that contained 10 mM phosphate buffer are shown in light colors and samples with 50 mM phosphate buffer are shown in dark colors. Squares represent GluC libraries, circles represent chymotrypsin libraries, and triangles represent trypsin libraries. (C) Fraction of peptide libraries labeled with 2PCA at 20 h over a range of temperatures from 37ºC to 75ºC. Samples that contained 10 mM phosphate buffer are shown in light colors and samples with 50 mM phosphate buffer are shown in dark colors. Squares represent GluC libraries, circles represent chymotrypsin libraries, and triangles represent trypsin libraries. (D) Fraction of peptide libraries labeled with 0.01-50 mM 2PCA. Squares represent GluC libraries, circles represent chymotrypsin libraries, and triangles represent trypsin libraries. (E) Residue-specific peptide library labeling of 2-PCA at concentrations from 0.01-50 mM. Data for individual libraries are shown in Fig. S5. Concentration-dependent specificity data are shown in Fig. S6.

We next sought to optimize the concentration of 2PCA for N-terminal modification (**Fig. 2D**). We measured an EC_50_ of 0.3 ± 0.1 mM for 2PCA modification of peptide libraries and found that the fraction of peptides modified increases as a function of 2PCA concentration below 10 mM. Increasing the 2PCA concentration above 10 mM did not substantially improve labeling. We also analyzed the concentration dependence of modification of peptides with a specific amino acid at position 1 or position 2 (**Fig. 2E, Fig. S5**). At position 1, we found that most N-terminal amino acids exhibit similar levels of modification at a given concentration of 2PCA. However, for peptides with Gly at their N termini, the fraction of peptides modified was lower than was observed for other N-terminal sequences at every 2PCA concentration. At position 2, peptides with any amino acid except for Pro and Gly showed similar levels of modification at a given 2PCA concentration. However, as expected, peptides with Pro at position 2 could not be modified at any concentration of 2PCA, while peptides with Gly at position 2 were modified somewhat more efficiently than other sequences. N-terminal specificity did not vary substantially across different concentrations of 2PCA (**Fig. S6**). Together, these experiments illustrate the parameters that affect the efficiency of 2PCA modification of complex samples. Our data show that maximizing the extent of 2PCA modification of a sample may require the use of high millimolar concentrations of the reagent. At the same time, much lower concentrations of 2PCA reagent used at physiological temperature on the timescale of hours are likely to be suitable for applications like enrichment N terminomics, in which quantitative modification of N termini is not required.

### Applying 2PCA reagents for ‘catch-and-release’ N terminomics

Biotinylated probes based on 2PCA have been used previously for proteome-wide N-terminal modification and enrichment for identification of protease cleavage sites (**Fig. 3A**)^7^. This method, termed <u>CH</u>emical enrichment <u>O</u>f <u>P</u>rotease <u>S</u>ubstrates (CHOPS) was applied to probe substrates of the dipeptidyl peptidases DPP8 and DPP9. The CHOPS method relies on biotin-2PCA (**2**) for N-terminal modification and enrichment, providing an effective strategy for isolating modified N termini. However, the high affinity between biotin and avidin presents difficulties for efficient recovery of biotinylated peptides. Additionally, biotin-2PCA is not commercially available and must be synthesized by the user. We sought to develop a chemically cleavable biotin-2PCA probe for one-step N-terminal biotinylation for N terminomics applications to enhance recovery of N-terminal peptides (**Fig. 3AB**). We also sought to eliminate the need for probe synthesis by integrating a commercially available, modular alkyne-2PCA probe with clickable, selectively cleavable biotin reagents for enrichment proteomics (**Fig. 3A, C**).

**Figure 3.**
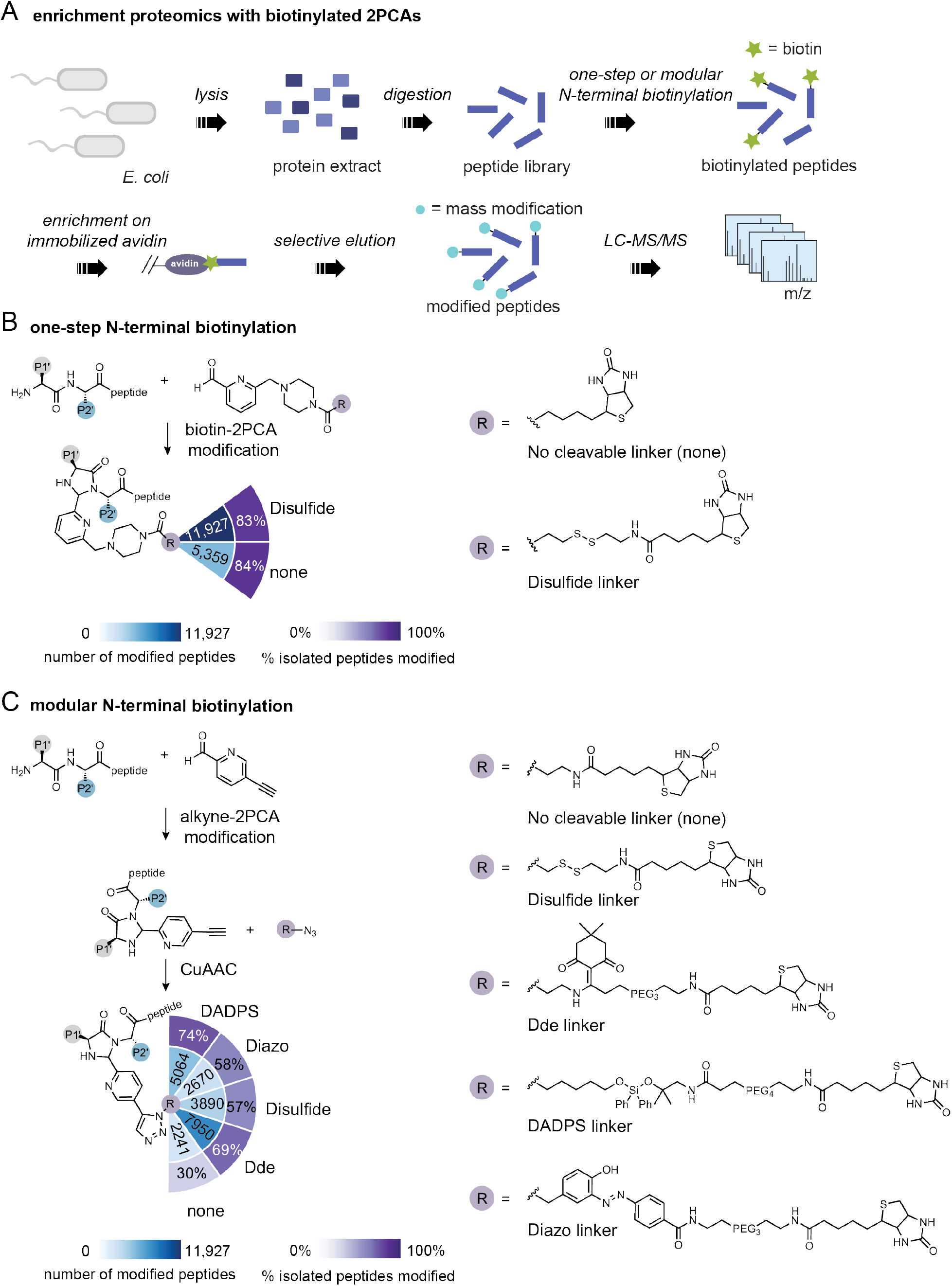
One-step or modular biotinylation with 2PCA reagents for ‘catch-and-release’ enrichment proteomics. (A) Workflow for N-terminal biotinylation of proteome-derived peptide libraries using biotinylated or clickable 2PCA reagents. Data showing the efficiency and specificity of alkyne-2PCA library modification are show in Fig. S7. (B) One-step N-terminal biotinylation of proteome-derived peptide libraries with biotin-2PCA or biotin-SS-2PCA followed by enrichment, elution, and LC-MS/MS analysis of modified peptides. (C) Modular N-terminal biotinylation of proteome-derived peptide libraries using alkyne-2PCA followed by CuAAC with chemically cleavable biotin azides, enrichment, selective elution, and LC-MS/MS analysis of modified peptides. Data showing the efficiency of the CuAAC reaction are shown in Fig. S8.

We first designed a biotin-disulfide-2PCA (biotin-SS-2PCA, **3**) probe to test whether use of a chemically cleavable linker would improve recovery of N-terminally biotinylated peptides (**Fig. 3B**). We then treated proteome-derived peptide libraries with either biotin-2PCA or biotin-SS-2PCA, enriched the peptides on immobilized neutravidin, and eluted the peptides by either heating the resin to 70ºC in 80% acetonitrile/0.1% formic acid for 10 min (for biotin-2PCA) or by treating the resin with 5 mM TCEP in 20 mM HEPES, pH 7.5 at room temperature for 1 h (for biotin-SS-2PCA) to cleave the disulfide linker. After analyzing the eluted peptides by LC-MS/MS, we found that although the fraction of recovered peptides that were 2PCA modified was similar across samples (84% for biotin-2PCA vs 83% for biotin-SS-2PCA), the number of 2PCA-modified peptides was more than twofold higher in the samples enriched with the chemically cleavable disulfide linker (5,359 peptides for biotin-2PCA vs. 11,927 for biotin-SS-2PCA). Chemically cleavable linkers can therefore be used to improve the efficiency of elution of biotinylated N-terminal peptides, as has been previously noted for other types of chemoproteomic probes^18–21^.

We next sought to develop a catch-and-release 2PCA probe based on commercially available reagents (**Fig. 3C**). We envisioned a strategy in which 2PCA with an alkyne substituent would be used to introduce a handle for click chemistry with cleavable biotin azides. Toward this goal, we tested the ability of 5-ethynylpicolinaldehyde (**4**, “alkyne-2PCA”) to modify N termini in proteome-derived peptide libraries. We found that alkyne-2PCA modified proteome-derived peptide libraries with similar efficiency and specificity to 2PCA (**Fig. S7**). We next tested whether alkyne-2PCA-modified peptides are substrates for copper (I)-catalyzed alkyne-azide cycloaddition (CuAAC)^22^ with biotin azide. We found that 95% of alkyne-2PCA peptides could be biotinylated in this manner (**Fig. S8**). Alkyne-2PCA modification followed by click chemistry with biotin azide therefore represents an efficient strategy to biotinylate peptide N termini using only commercially available reagents without the need to synthesize and purify a biotin-2PCA probe.

Numerous chemically cleavable biotin azides are commercially available. We set out to determine which chemically cleavable linker produced the highest recovery of N-terminally biotinylated peptides (**Fig. 3C**). We compared biotin azide (**5**), biotin-SS-azide (**6**), biotin-Dde-azide (**7**), biotin-DADPS-azide (**8**), and biotin-Diazo-azide (**9**) by performing click chemistry under identical conditions, enriching the biotinylated peptides, and then selectively eluting the captured peptides under conditions specific to each linker (as described in **Methods**). We found that all four chemically cleavable linkers outperformed biotin azide in terms of the number of recovered peptides that were identified in LC-MS/MS experiments. The largest number of peptides were identified in samples enriched with biotin-Dde-azide (7,950 peptides), followed by biotin-DADPS-azide (5,064 peptides), biotin-disulfide-azide (3,890 peptides), biotin-Diazo-azide (2,670 peptides), and biotin azide (2,241 peptides). Alkyne-2PCA modification combined with clickable, cleavable biotin azides is therefore an off-the-shelf system for N-terminal modification and enrichment.

### Proteomic Identification of Cleavage Sites with 2PCA reagents (PICS2)

We next set out to apply 2PCA-based catch-and-release reagents for protease sequence specificity profiling (**Fig. 4**). Based on their diversity and ease of preparation, proteome-derived peptide libraries are useful as pools of substrates for profiling protease specificity^3^. However, most existing methods for isolating proteolytic neo-N termini from proteome-derived peptide libraries do not provide a straightforward means of characterizing proteases that recognize Lys in their consensus cleavage sequences. For example, the widely adopted PICS (Proteomic Identification of Cleavage Sites) method for protease specificity profiling in proteome-derived peptide libraries requires blocking of N termini and Lys side chains prior to protease digestion to enable biotinylation of proteolytic neo-N termini with an amine-reactive reagent^3^. This blocking step precludes the use of PICS for characterization of biologically critical proteases with Lys in their consensus cleavage sequences, including those involved in cancer^23–25^, inflammation^25^, and viral infection^26–28^. We sought to leverage the N-terminally selective reaction of 2PCA reagents to circumvent these limitations. We envisioned a workflow in which N termini would be selectively blocked, leaving lysine ε-amines unblocked. Following treatment with a test protease of interest, proteolytic neo-N termini would be selectively modified with biotin-SS-2PCA for enrichment on neutravidin resin and subsequent identification using LC-MS/MS. We termed this strategy <u>P</u>roteomic Identification of protease <u>C</u>leavage <u>S</u>ites with <u>2</u>PCA reagents (PICS2) (**Fig. 4A**).

**Figure 4.**
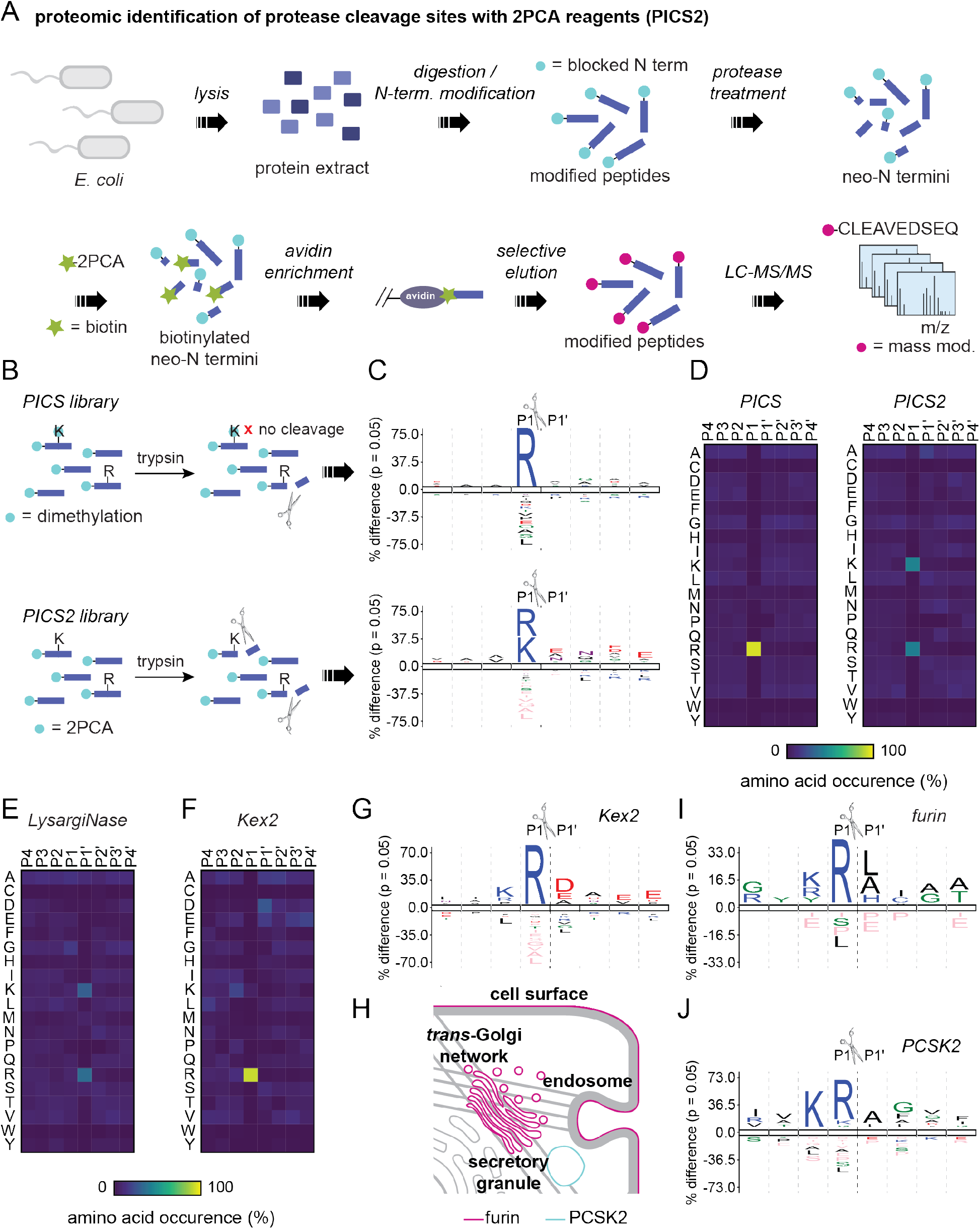
Proteomic identification of protease cleavage sites with 2PCA reagents (PICS2). (A) Proteomic identification of protease cleavage sites with 2PCA reagents workflow. Proteome-derived peptide libraries are N-terminally blocked prior to protease treatment. After treatment with the protease of interest, neo-N termini are selectively modified with biotin-SS-2PCA, enriched, selectively eluted, and analyzed by LC-MS/MS. IceLogos for human proteome-derived peptide libraries are shown in Fig. S9. Data showing the efficiency of N-terminal blocking with 2PCA are shown in Fig. S10. (B) Comparison of PICS and PICS2 libraries and expected trypsin cleavage specificity. (C) IceLogos for trypsin cleavage specificity determined by PICS (top) or PICS2 (bottom). (D) Heatmaps showing the percent amino acid occurrence in each of the peptide subsites P4-P4‘ for trypsin PICS (left) and PICS2 (right). Results for individual proteome-derived peptide libraries are shown in Fig. S11. (E) Heatmap showing the percent amino acid occurrence in each of the peptide subsites P4-P4‘ for LysargiNase PICS2. Results for individual proteome-derived peptide libraries are shown in Fig. S12. Additional data demonstrating the PICS2 method for GluC and chymotrypsin is shown in Fig. S13 and Fig. S14. (F) Heatmap showing the percent amino acid occurrence in each of the peptide subsites P4-P4‘ for Kex2 PICS2. Data showing the efficiency and specificity of N-terminal blocking by dimethylation is shown in Fig. S15. Kex2 results with individual peptide libraries are shown in Fig. S16. Previously known Kex2 substrates are shown in Fig. S17. (G) IceLogo for Kex2 cleavage specificity as determined by PICS2. (H) Subcellular localization of furin and PCSK2. (I) IceLogo for furin cleavage specificity as determined by PICS2. Furin PICS2 results with individual peptide libraries are shown in Fig. S18. (J) IceLogo for PCSK2 cleavage specificity as determined by PICS2. PCSK2 PICS2 results with individual peptide libraries are shown in Fig. S19.

To test the PICS2 strategy, we generated proteome-derived peptide libraries by digesting *E. coli* or human lysates with either chymotrypsin (P1 = F, L, W, or Y) or GluC (P1 = D or E) (**Fig. S1, Fig. S9**). After isolating digested peptides, the libraries were N-terminally blocked using 2PCA. LC-MS/MS analysis demonstrated that 89±0.2% of peptides in the human library and 81±5% of peptides in the *E. coli* library were N-terminally modified by 2PCA under these conditions (**Fig. S10**). We first benchmarked the PICS2 workflow using trypsin, a well characterized protease that cleaves after lysine and arginine. As a comparison point, we also analyzed trypsin specificity using PICS (**Fig. 4BCD, Fig. S11**)^3^. In total we identified 2,984 neo-N termini from the PICS workflow and 1,011 neo-N termini from the PICS2 workflow. Although both methods enabled enrichment of >1,000 tryptic neo-N termini, the tryptic peptides identified in the PICS experiment were limited to those with Arg at the P1 position based on the inability of trypsin to cleave after dimethylated lysine residues (**Fig. 4B**). In contrast, PICS2 enabled identification of neo-N termini derived from sequences containing both Lys and Arg in the P1 position, more accurately reflecting trypsin’s well characterized specificity (**Fig. 4CD**).

To test the generality of the PICS2 method, we used the workflow to characterize the specificity of a panel of commercial proteases comprised of LysargiNase29 (P1‘ = K or R) (**Fig. 4E, Fig. S12**), GluC (P1 = D or E) (**Fig. S13**), and chymotrypsin (P1 = F, L, W, or Y) (**Fig. S14**). We found that the specificity profiles generated by PICS2 analysis matched the known specificity of each protease.

We also tested a previously reported method for blocking peptide N termini prior to protease digestion, reductive dimethylation under acidic conditions30. We found that this strategy blocked N termini similarly to 2PCA (89%±9% vs 89%±0.2% for 2PCA) (**Fig. S15**), but also resulted in modification of 14%±4% of peptides on Lys residues. This strategy is not ideal for biotinylating proteolytic neo-N termini for enrichment because it is expected to result in substantial Lys labeling as well as double modification of the N terminus, necessitating higher reagent concentrations and more stringent elution conditions. However, reductive dimethylation has the advantage that it requires only 10 min of reaction time while sill leaving most Lys residues unmodified. Therefore, combining N-terminally selective reductive dimethylation for N-terminal blocking and 2PCA-based biotinylation for N-terminal capture represents an alternative strategy for protease specificity profiling with proteome-derived peptide libraries.

### Profiling the sequence specificity of proprotein convertases with PICS2

Subtilisin/kexin-type proprotein convertases (PCSKs) are an important class of eukaryotic serine proteases that are involved in numerous (patho)physiological processes^9,10^. PCSKs are expressed in the secretory pathway and cleave protein and peptide precursors, including those of peptide hormones, enzymes, and cell surface receptors, to generate their mature, active forms. In humans, there are nine PCSK family members: PCSK1, PCSK2, furin, PCSK4, PCSK5, PCSK6 (also known as PACE4), PCSK7, SKI-1, and PCSK9. The first seven of these cleave proprotein substrates on the C-terminal side of single or paired basic amino acid residues, while SKI-1 cleaves following non-basic residues and PCSK9 cleaves itself following a Gln residue and is its own sole proteolytic substrate10. Based on their roles in human diseases, including hypercholesterolemia, cancer, diabetes, osteoarthritis, and viral and bacterial infection, PCSKs represent attractive drug targets^9,10^. However, because of their crucial physiological functions and overlapping substrate specificity, development of small-molecule inhibitors of individual PCSKs remains challenging. A detailed view of PCSK sequence specificity could therefore reveal differences in molecular recognition between PCSK family members that could inform inhibitor design. We therefore sought to apply PICS2 for PCSK sequence specificity profiling.

We initially examined *S. cerevisiae* Kex2, a well-studied PCSK that typifies the family (**Fig. 4FG, Fig. S16**). Kex2 has two well-characterized physiological substrates: the yeast mating pheromone pro-α-factor and the secreted pore-forming protein K1 killer toxin^31–33^. Pro-α-factor is cleaved at Lys-Arg85, Lys-Arg104, Lys-Arg125, and Lys-Arg146, while K1 killer toxin is cleaved at Pro-Arg44, Arg-Arg149, Lys-Arg188, and Lys-Arg233. Although active Kex2 is required for all four killer toxin cleavage events, it has remained unclear whether Kex2 or a second Kex2-dependent protease cleaves at the Pro-Arg site33. PICS2 analysis of Kex2 revealed that Kex2 has a stringent preference for Arg at P1, but accepts Lys, Arg, or Pro at P2 (**Fig. 4FG**), supporting the hypothesis that Kex2 directly cleaves the Pro-Arg site in K1 killer toxin. PICS2 analysis also revealed that Kex2 prefers acidic residues on the prime side of the cleavage site, particularly at the P1‘, P3‘, and P4‘ positions. This result is consistent with the prime side sequences of the cleavage sites in pro-α-factor (P1‘-<u>E</u>A<u>E</u>A-P4‘ or P1‘-<u>E</u>A<u>D</u>A-P4‘) and K1 killer toxin (P1‘-<u>E</u>APW-P4‘, P1‘-<u>D</u>IST-P4‘, P1‘-S<u>D</u>TA-P4‘, and P1‘-YVYP-P4‘) (**Fig. S17**).

We next turned our attention to the human PCSKs. Human PCSK substrate specificity is largely determined by subcellular localization. While PCSK1 and PCSK2 are restricted to acidic regulated secretory granules^10,34^, furin and the other PCSKs are localized to the *trans*-Golgi network (TGN), cell surface, extracellular matrix, and/or endosomes (**Fig. 4H**)^10,35^. To analyze the specificity of human PCSKs, we generated human proteome-derived peptide libraries from HEK293T cell protein extracts (**Fig. S9**). We used these libraries to profile the specificity of PCSK2, a secretory granule-localized proprotein convertase, and furin, which resides in the TGN, at the cell surface, and in endosomes. We found that furin has a stringent preference for Arg at P1, but will accept Lys, Arg, or Tyr at P2 (**Fig. 4I, Fig. S18**). At P4, furin prefers Arg or Gly. In contrast, PCSK2 accepts both Lys and Arg at P1, but strongly prefers Lys at P2 (**Fig. 4J, Fig. S19**). At P4, PCSK2 prefers Arg, Ile, or Val. These results reveal that despite their generally similar substrate sequence specificity, furin and PCSK2 each have unique sequence recognition capacities that could potentially be exploited for selective inhibitor design.

### Mapping the apoptotic proteome with 2PCA catch-and-release reagents

While proteome-derived peptide library-based methods can reveal protease sequence specificity, understanding cellular proteolytic signaling pathways also requires identification of protease substrates that are cleaved in living cells in response to a stimulus of interest. Global sequencing of cellular N termini, or ‘N terminomics’, can provide key insights into how proteins are modified by proteolytic cleavage to alter their biological functions^4,5,7,36^. Numerous methods that exploit the chemical properties of N termini, including their charge, reactivity, and structure, have been developed to enable global analysis of both proteolytic and translational N termini using mass spectrometry-based proteomics^4–7,37^. Positive enrichment methods for N terminomics rely on specific modification of protein N termini with an affinity handle to directly isolate them from complex samples such as cell lysates^4,7^, while other methods depend on depletion of internal peptides following trypsin digestion^5,37^. Positive enrichment approaches have the advantage of high sensitivity for low-abundance peptides but present the challenge that N-terminal α-amines must be selectively modified over lysine ε-amines. Existing methods for positive enrichment of protein N termini include subtiligase N terminomics^4,11,38,39^, which employs the designed peptide ligase subtiligase for selective N-terminal biotinylation, and CHOPS7, which uses biotin-2PCA for N-terminal modification. Subtiligase N terminomics has the advantage that it uses a catch-and-release strategy in which biotinylated N-terminal peptides can be selectively eluted from immobilized avidin using a TEV protease cleavage site built into the biotinylated subtiligase substrates^4^. However, the method is limited by sequence specificity inherent to subtiligase^11^ and the need to synthesize a specialized biotinylated peptide ester probe^40^. In contrast, CHOPS has little sequence specificity, but existing workflows require chemical synthesis of the biotin-2PCA reagent and do not offer an approach for N-terminal catch-and-release7.

We hypothesized that commercially available 2PCA reagents could be used to implement a catch-and-release N terminomics strategy, simultaneously overcoming the limitations of both subtiligase N terminomics and CHOPS. We refer to this approach as CHOPPER (<u>Ch</u>emical enrichment <u>O</u>f <u>P</u>rotease substrates with <u>P</u>urchasable, <u>E</u>lutable <u>R</u>eagents) (**Fig. 5**). We sought to apply this approach to map proteolysis during apoptosis, a programmed cell death pathway executed by the caspases, a family of cysteine proteases that cleave C-terminal to aspartate residues (P1 = D)^41,42^. During apoptosis, specific caspase cleavages in hundreds of proteins across the proteome lead to changes in protein function that program membrane blebbing, cytoskeletal rearrangements, chromatin condensation, DNA fragmentation, and ultimately, destruction of the dying cell^43^. Apoptosis has been studied extensively with N terminomics, providing an opportunity to benchmark our approach. However, the limitations of existing N terminomics methods also offer an opportunity for biological discovery with CHOPPER.

**Figure 5.**
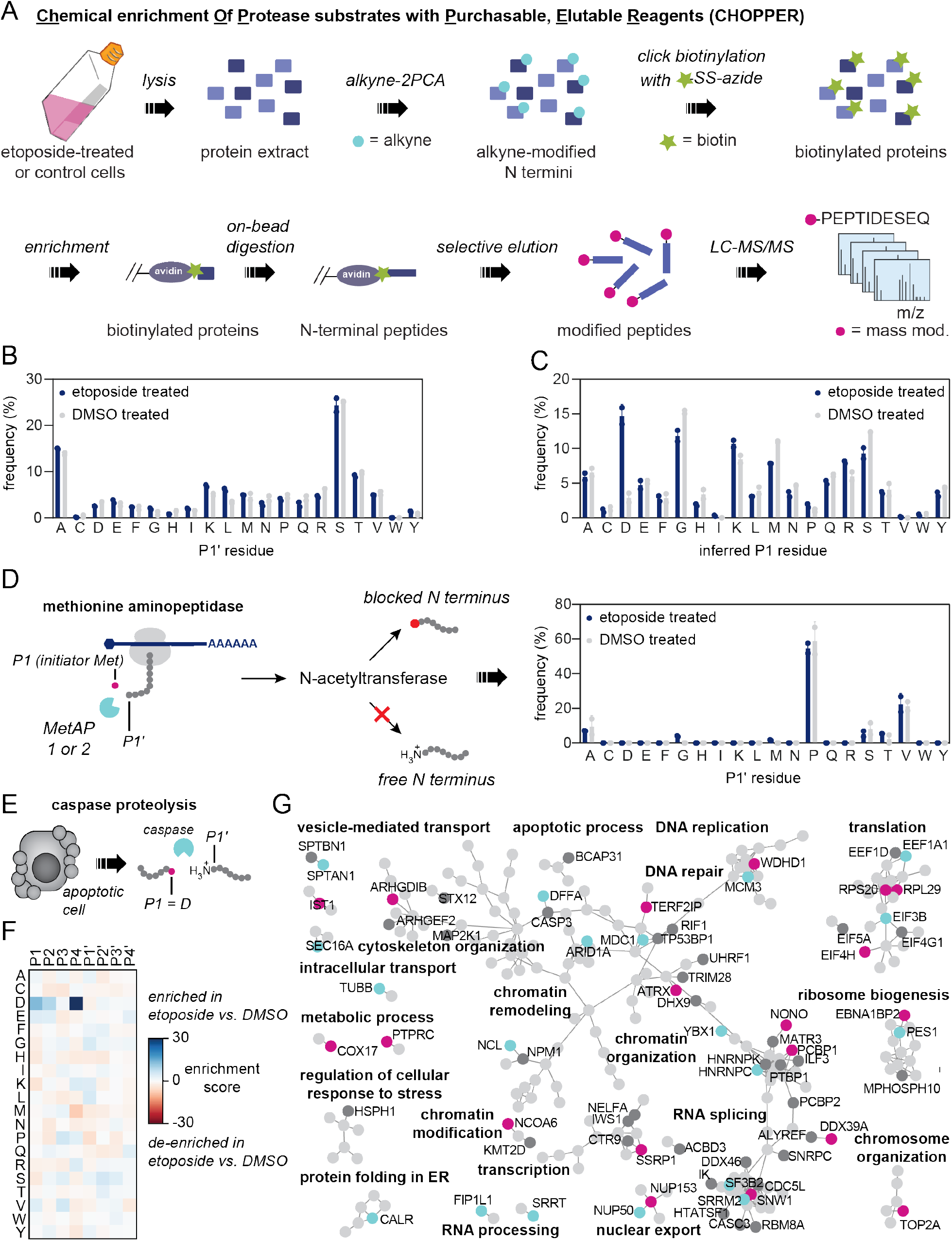
Chemical enrichment of protease substrates with purchasable, elutable reagents (CHOPPER). (A) Analysis of apoptotic proteolysis with CHOPPER. Jurkat cells are treated with etoposide or DMSO vehicle and lysed. The lysate is modified with alkyne-2PCA followed by CuAAC with biotin-SS-azide. Biotinylated proteins are enriched and digested on bead. N-terminal peptides are selectively eluted by cleavage of the disulfide bond and analyzed by LC-MS/MS. Images of etoposide- and DMSO-treated cells are shown in Fig. S20. (B) Frequency of each amino acid in the P1‘ position for etoposide-treated and DMSO-treated cells. (C) Frequency of each amino acid at the inferred P1 position for etoposide-treated and DMSO-treated cells. (D) Frequency of each amino acid at the P1’ position of methionine aminopeptidase cleaved, unacetylated substrates. (E) Caspase proteolysis is characterized by P1 = D. (F) Heatmap showing enrichment or de-enrichment of each amino in each position among N termini enriched from etoposide-treated cells vs. DMSO-treated cells. (G) STRING analysis of putative caspase cleavages identified in the CHOPPER dataset and in subtiligase N terminomics datasets. Light grey circles represent proteins that were previously known to be caspase substrates, dark grey circles represent proteins that were previously known to be caspase substrates and were also identified by CHOPPER, blue circles represent proteins in which caspase cleavage sites were previously known but a new cleavage site was identified by CHOPPER, and magenta circles represent proteins that were not previously known to be caspase substrates in etoposide-treated Jurkat cells. Full STRING analysis is shown in Fig. S21. Positions of putative caspase cleavages relative to protein domain boundaries are shown in Fig. S22. A comparison between N termini captured with CHOPPER and subtiligase N terminomics is shown in Fig. S23.

To test this strategy, we generated lysates from Jurkat cells treated with the apoptosis inducer etoposide or treated with a vehicle control (**Fig. 5A**). After 8 h, etoposide-treated cells exhibited extensive blebbing characteristic of apoptosis and were harvested for analysis (**Fig. S20**). Lysates were treated with 1 mM alkyne-2PCA and then subjected to click chemistry with biotin-disulfide-azide. Proteins with biotinylated N termini were then enriched on neutravidin resin and digested with trypsin. N-terminal peptides were selectively eluted by reduction of the disulfide bond and analyzed by LC-MS/MS. In total, we identified 1587 unique N-terminal peptides in the apoptotic samples and 654 unique N-terminal peptides in the vehicle control samples. Across all peptides in the samples, no unexpected biases at the 2PCA-modified P1‘ position were observed, reflecting the broad sequence compatibility of 2PCA reagents (**Fig. 5B**). However, consistent with induction of caspase activity (P1 = D), we observed a substantial increase in the fraction of peptides with P1 = D in the etoposide-treated samples (**Fig. 5C**).

To benchmark CHOPPER, we first analyzed substrates of the methionine aminopeptidases (MetAPs), whose activity is not known to be affected by apoptosis, in both apoptotic and nonapoptotic samples (**Fig. 5D**). In human cells, two MetAPs, MetAP1 and MetAP2, are involved in co-translational cleavage of Met1 from proteins. Previous biochemical studies have shown that human MetAPs favor Ala, Cys, Gly, Pro, Val, and Ser at the P1‘ position44. Following Met excision, substrates with N-terminal Ala, Cys, Gly, Val, or Ser can be further modified by the N-acetyltransferase (NAT) NatA, resulting in a blocked N terminus inaccessible to 2PCA modification. Six other human NATs modify other N-terminal sequences, and it is estimated that >85% of cytosolic proteins are N-terminally acetylated. However, no human NAT is known that acetylates proteins with N-terminal Pro45. Across our apoptotic and non-apoptotic samples, we identified a total of 73 putative MetAP substrates based on the presence of the 2PCA tag at residue 2 (**Fig. 5D**). We did not observe a substantial difference in the distribution of P1‘ residues captured in apoptotic versus non-apoptotic samples. Interestingly, >50% of putative MetAP substrates in all samples had P1‘ = P. While this distribution is likely consistent with the *in vivo* balance of MetAP activity and NAT activity45, this result differs from a previous examination of MetAP susbtrates using subtiligase N terminomics in which 0-2% of identified MetAP substrates had P1‘ = P11. This difference likely arises from subtiligase substrate specificity, which is strongly biased against P1‘ = P, highlighting the complementary utility of CHOPPER.

We next analyzed putative caspase substrates with P1 = D in our datasets (**Fig. 5E**). While nonapoptotic samples had 21/654 (3%) P1 = D peptides, 229/1587 (14%) N-terminal peptides derived from etoposide-treated cells had P1 = D, a clear enrichment consistent with induction of caspase activity (**Fig. 5C,F**). We compared our datasets to two previously published subtiligase N terminomics datasets that examined etoposide-treated Jurkat cells: one published in Mahrus et al.^4^ and deposited in the Degrabase (298 unique P1 = D sites)^46^ and one published in Weeks and Wells, 2018 (1,152 P1 = D sites)^11^. We found that 112/229 cleavage events in 95 proteins had not been previously reported, including cleavages in 44 proteins that were not previously identified as substrates of etoposide-induced caspase activity in Jurkat cells (**Table S1**). STRING analysis of cleavages newly identified by CHOPPER and previously reported etoposide-induced caspase cleavages demonstrated that the new cleavage events largely affect the same pathways that are known to be targeted by caspases during apoptosis (**Fig. 5G, Fig. S21**). These include RNA splicing, DNA repair, transcription, translation, and intracellular transport, among others^4^. Many of the newly identified cleavages occur between domain boundaries of multidomain proteins (**Fig. S22**). Notably, many of these substrate proteins contain domains that modulate protein-protein interactions (e.g., SH3, PDZ, BRCT, FHA, and ADD domains), protein-DNA interactions (e.g., Myb-like and bHLH domains), or protein-RNA interactions (e.g., KH and RRM domains)^47^, suggesting that apoptotic proteolysis leads to disruption or reprogramming of the spatial organization of macromolecular complexes. This observation is consistent with numerous previous studies of specific caspase cleavages^4,48–51^.

To understand the features of the CHOPPER workflow that enabled identification of additional caspase cleavage sites, we examined the distribution of amino acids at the P1‘ and P2‘ positions of P1 = D peptides (**Fig. S23**). Compared to subtiligase or a cocktail of subtiligase specificity mutants^11^, CHOPPER performed better on peptides with branched-chain amino acids or Pro at the P1‘ position. This difference is likely based on the substrate specificity of subtiligase, for which peptides with branched-chain amino acids or Pro in the first position are poor substrates. CHOPPER also identified a higher fraction of peptides with Lys or Arg at P1‘. While subtiligase can efficiently modify substrates with Lys or Arg at P1‘, subsequent trypsin digestion is expected to cleave the subtiligase biotin modification from the peptide. We hypothesize that in the CHOPPER workflow, cyclization of the N-terminal Lys or Arg inhibits cleavage by trypsin, enabling enrichment of P1‘ = Lys or Arg peptides. In contrast, subtiligase performs better for substrates with P1‘ = Gly, consistent with the low modification efficiency for P1‘ = Gly observed for 2PCA modification of proteome-derived peptide libraries. At P2‘, CHOPPER outperforms subtiligase for peptides with Gly or Ser in this position, while subtiligase performs better for peptides with branched-chain amino acids in the second position. These results are consistent with subtiligase’s preference for aromatic or large hydrophobic amino acids at P2‘. CHOPPER also captures a larger percentage of P2‘ = Lys or Arg substrates, suggesting that cyclization of the N-terminal amino acid may also inhibit trypsin cleavage C-terminal to nearby amino acid. Together, these results demonstrate that CHOPPER is a complementary technology to existing positive-enrichment N terminomics methods that enables more comprehensive proteolysis mapping in cell lysates.

## Discussion

Site-specific chemoproteomic and chemoenzymatic probes are valuable tools for analyzing the properties of proteins, including their activity, localization, and post-translational modification state. Probes exist to modify at least nine amino acids side chains and protein N and C termini with varying degrees of efficiency and specificity^16^. The utility of any probe depends on at least three factors: 1) predictable site and sequence specificity; 2) optimized modification conditions for proteomics applications; and 3) their accessibility to the research community. The expanded 2PCA-based toolbox that we report here addresses these three features through deep profiling of 2PCA specificity, optimization of reaction conditions for modification of proteomic samples, and incorporation of commercially available reagents that remove the need for chemical synthesis of the probes.

The use of proteome-derived peptide libraries for deep profiling of 2PCA specificity overcomes the challenges and limitations associated with other specificity profiling methods. Small synthetic peptide libraries cannot capture how amino acid identity at one position influences modification efficiency at another. Peptide libraries displayed on yeast^52,53^ or phage^54–56^ provide high sequence diversity but require a different set of conditions and assays that may not be directly relevant to proteomic samples and do not provide single amino acid resolution. Proteome-derived peptide libraries strike a balance, providing a high level of diversity that can be measured with the same LC-MS/MS assays and modified under the same conditions as proteomic samples^3,11^. Using proteome-derived peptide libraries, we found that peptides with Gly in the first position are generally modified with lower efficiency than other sequences, but that this effect can be overcome when certain other residues are present in the second position. This kind of positional cooperativity is common in enzymes but is often overlooked with chemical probes. We also found that, contrary to results from experiments in which only the N-terminal amino acid of a defined peptide sequence was varied^12^, 2PCA can efficiently modify peptides with Pro in the first position. This suggests that the low efficiency of P1‘ Pro modification may have been related to a specific feature of the peptide sequence that was used rather than an inherent bias of the 2PCA reagent. Importantly, the results of proteome-derived peptide library specificity experiments directly translated to the ability of 2PCA to modify protein N termini in cell lysates. For example, the ability of 2PCA to modify P1‘ Pro sequences enabled us to identify a larger number of human methionine aminopeptidase substrates than had been identified in previous enrichment N terminomics experiments^11^. Additionally, 2PCA specificity profiling in proteome-derived peptide libraries explains why 2PCA probes can capture N-terminal sequences that were not identified by other methods. Finally, by profiling 2PCA specificity using proteome-derived peptide libraries, we confirmed that 2PCA is unable to modify substrates with P2‘ Pro regardless of the identity of amino acids in other positions. This result translated to cell lysate samples and represents the only major sequence bias of 2PCA N-terminal modification.

Optimization of 2PCA modification using proteome-derived peptide libraries enabled us to identify conditions for selective, near-complete blocking of peptide N termini and conditions that balance a high level of N-terminal modification with use of low millimolar concentrations of biotinylated or alkyne-modified 2PCA reagent. Both insights were important for implementation of PICS2, a protease specificity profiling method that is compatible with proteases containing Lys or Arg in their cleavage sequences. High concentrations of 2PCA produced the N-terminally blocked proteome-derived peptide libraries that are needed for PICS2, while low millimolar concentrations of 2PCA reagents were sufficient for N-terminal alkyne modification and subsequent click chemistry with biotin and capture on neutravidin resin while avoiding competition from excess reagent. Similarly, low millimolar alkyne-2PCA concentrations were used for CHOPPER profiling of apoptotic protease substrates. The incorporation of a chemically cleavable linker between 2PCA and biotin enhances the efficiency of N-terminal peptide elution, providing more comprehensive substrate and specificity profiling and lowering sample requirements.

Incorporation of alkyne-2PCA as a modular means of N-terminal biotinylation improves accessibility of both PICS2 and CHOPPER to protease researchers. Probes in which 2PCA and biotin are covalently linked must be chemically synthesized and purified before use. Although 6- (1-piperazinylmethyl)-2-pyridinecarboxaldehyde12 recently became commercially available, reducing the synthesis to one step, synthesis and purification nonetheless create a barrier to the use of 2PCA reagent for N terminomics applications. In contrast, alkyne-2PCA and a variety of chemically cleavable biotin azides are commercially available, removing the need for chemical synthesis and purification and making the tools accessible to researchers without the specialized equipment and expertise needed for chemical synthesis. Although beyond the scope of this study, alkyne-2PCA is also likely to be useful in N-terminal bioconjugation to proteins, where it can provide a modular means of attaching any azide-modified payload to the protein N terminus.

The PICS2 and CHOPPER workflows are widely applicable for studying the substrates and specificity of a broad array of proteases. These workflows have well defined specificity with few limitations and are easy to implement using commercially available reagents. We anticipate that the expanded 2PCA-based N terminomics toolbox can be widely adopted based on its N-terminal selectivity, broad sequence compatibility, optimized reaction conditions, and ease of use, and will propel discovery in broad areas of protease research.

## Acknowledgements

We thank S. Coyle, L. Mazurkiewicz, K. Radziwon, M. Ravalin, and D. Sashital for helpful discussions. This work was supported in part by startup funds from the University of Wisconsin-Madison Department of Biochemistry, by a David and Lucille Packard Fellowship for Science and Engineering, and by a Career Award at the Scientific Interface from the Burroughs Wellcome Fund (1017065) (to A.M.W.). H.N.B was supported in part by a William R. and Dorothy E. Sullivan Wisconsin Distinguished Graduate Fellowship. C.L.F. was supported in part by the UW-Madison Biotechnology Training Program under grant number NIH 5 T32 GM135066.

## Author contributions

A.M.W. and H.N.B. designed the experiments. H.N.B., C.L.F., and A.M.W. performed experiments. A.M.W. and H.N.B. analyzed data and wrote the manuscript.

## Methods

### Key chemicals and materials

2-Pyridinecarboxaldehyde (**1**) and 6-(1-piperazinylmethyl)-2-pyridinecarboxaldehyde bistosylate salt were purchased from Sigma-Aldrich. 5-Ethynylpicolinaldehyde (“alkyne-2PCA”) (**4**) was purchased from Ambeed. NHS-biotin, NHS-SS-biotin, SOLA HRP SPE cartridges, Pierce Snap-Cap spin columns, and High-Capacity Neutravidin Agarose were purchased from Thermo Fisher Scientific. Biotin azide (**5**) was purchased from Cayman Chemical. Azide-SS-biotin (**6**) was purchased from Broadpharm. Biotin-Diazo-azide (**9**), biotin-Dde-azide (**7**), and biotin-DADPS-azide (**8**) were purchased from Click Chemistry Tools. Full chemical structures of key reagents are shown in **Fig. S24**.

### Generation of *E. coli* proteome-derived peptide libraries

LB media (1 L) was inoculated with a single colony of *E. coli* XL10. Cells were grown overnight at 37ºC with shaking at 200 rpm. Cells were harvested by centrifugation at 4,000 × g for 10 min at 4ºC. The cell pellet was resuspended in 50 mL of lysis buffer (10 mM HEPES, 1 mM PMSF, 0.5 mM EDTA) and cells were lysed by three passes through a homogenizer at 15,000 psi. Cell debris was pelleted by centrifugation at 10,000 × g for 20 min at 4ºC. The supernatant was adjusted to 100 mM HEPES, pH 7.5. DTT was added to a final concentration of 5 mM and the lysate was incubated for 1 h at room temperature. Iodoacetamide was then added to a final concentration of 10 mM and the sample was incubated in the dark for 1 h at room temperature. Protein was precipitated by addition of trichloroacetic acid (TCA) to 15% (w/v) followed by a 2-6 h incubation at -20ºC. Proteome-derived peptide libraries were then produced as described previously.

### Generation of human proteome-derived peptide libraries

HEK293T cells (ATCC #CRL-3216) were grown to 90% confluence in DMEM media supplemented with 10% (v/v) fetal bovine serum at 37ºC under a 5% CO2 atmosphere. Cells were washed once with PBS and versene (10 mL) was added. The flask was incubated for 10 min at 37ºC. Cells were resuspended in versene, transferred to a conical tube, and harvested by centrifugation at 300 × g for 5 min. The cell pellet was resuspended in 800 μL hot lysis buffer (100 mM Tris-HCl, pH 8.5, 6 M guanidine hydrochloride, 5 mM TCEP, 10 mM chloroacetamide) that had been preheated to 95ºC and the suspension was heated at 95ºC for 10 min. Lysis was completed using 10 cycles of probe ultrasonication (20% amplitude; 5 s on / 5 s off). Insoluble material was removed by centrifugation at 20,000 × g at 4ºC for 10 min. Protein was then precipitated by addition of TCA to a final concentration of 15% (w/v). Proteome-derived peptide libraries were then produced as described previously. HEK293T cells used for proteome-derived peptide library generation were tested every six months for mycoplasma contamination using the LookOut Mycoplasma PCR Detection Kit (Sigma-Aldrich) according to the manufacturer’s instructions (**Fig. S25**).

### Modification of proteome-derived peptide libraries with 2PCA

*E. coli* or human proteome-derived peptide library modification reactions contained 1 mg/mL proteome-derived peptide library, the appropriate concentration of 2-PCA (10 μM-50 mM), and phosphate buffer, pH 7.5 (10 mM or 50 mM). Temperature dependence reactions contained 50 mM 2PCA and were incubated at 37ºC-75ºC for the indicated amount of time (4 h or 20 h). For 2PCA specificity profiling (**Fig. 1**), reactions contained 1 mg/mL peptide library, 10 mM 2PCA, and 50 mM sodium phosphate, pH 7.5 and were incubated for 4 h at 37ºC. For N-terminal blocking of peptide libraries, reactions contained 1 mg/mL peptide library, 50 mM 2PCA, and 50 mM sodium phosphate, pH 7.5 and were incubated for 20 h at 37ºC. After 2PCA modification, samples were desalted using the single-pot, solid-phase-enhanced sample-preparation (SP3) protocol57. Beads were prepared by mixing hydrophilic and hydrophobic Sera-Mag speedbeads (Cytiva Cat. #45152105050250 and Cat. #65152105050250) in a 1:1 ratio. Tubes were placed on a magnetic stand and supernatants were removed. Bead mixtures were then washed three times with water (1 mL). Supernatants were removed and tubes were removed from the stand. Acetonitrile was added to peptide samples to a final concentration of 95% (v/v) and samples were transferred to tubes containing the prewashed beads in ratio of 10 μg beads to 1 μg peptide. Tubes were vortexed briefly and then incubated at 30ºC on a Thermomixer (Eppendorf) at 1000 rpm for 10 min. Samples were placed on a magnetic stand and supernatant was removed. Beads were then washed three times with 1 mL acetonitrile. After the final wash, beads were air-dried for two minutes. A volume of water equivalent to ten times the original bead volume was added for elution. Beads were sonicated for 1 min in a water bath sonicator and then incubated at 30ºC for 5 min on a Thermomixer at 1000 rpm. Tubes were placed on a magnetic stand and eluted peptides were collected by pipetting into a fresh tube.

### Synthesis of biotin-2PCA

Biotin-2PCA (**2**) was synthesized as described previously^12^. To a solution of 6-(1-Piperazinylmethyl)-2-pyridinecarboxaldehyde bistosylate salt (25 mg, 0.045 mmol, 1 equiv.) in *N, N*-dimethylformamide (DMF) was added triethylamine (13.6 mg, 18.8 μL, 0.135 mmol, 3 equiv.) and NHS-biotin (17 mg, 0.05 mmol, 1.1 equiv.). The reaction mixture was stirred for 1 h at room temperature. The mixture was injected onto an Agilent Eclipse XDB-C18 column (9.4 × 250 mm, 5 μm) and purified by reverse-phase HPLC (0-100% B over 90 min at 2 mL/min; A: 0.1% aqueous TFA, B: 0.1% TFA in acetonitrile) using an Agilent 1260 quaternary pump coupled to a diode-array detector. Fractions (1 mL) were flash frozen in liquid nitrogen and lyophilized. The lyophilized biotin-2PCA was dissolved in DMSO and characterized using an Agilent G6230B time-of-flight mass spectrometer (TOF-MS) (m/z (M+H^+^), 432.2081; m/z (M+H^+^_calc_), 432.2069).

### Synthesis of biotin-SS-2PCA

Biotin-SS-2PCA (**3**) was synthesized using the general protocol described previously^12^. To a solution of 6-(1-Piperazinylmethyl)-2-pyridinecarboxaldehyde bistosylate salt (25 mg, 0.045 mmol, 1 equiv.) in *N, N*-dimethylformamide (DMF) was added triethylamine (13.6 mg, 18.8 μL, 0.135 mmol, 3 equiv.) and NHS-SS-biotin (25 mg, 0.05 mmol, 1.1 equiv.). The reaction mixture was stirred for 1 h at room temperature. The mixture was injected onto an Agilent Eclipse XDB-C18 column (9.4 × 250 mm, 5 μm) and purified by reverse-phase HPLC (0-100% B over 90 min at 2 mL/min; A: 0.1% aqueous TFA, B: 0.1% TFA in acetonitrile) using an Agilent 1260 quaternary pump coupled to a diode-array detector. Fractions (1 mL) were flash frozen in liquid nitrogen and lyophilized. The lyophilized biotin-SS-2PCA was dissolved in DMSO and characterized using an Agilent G6230B time-of-flight mass spectrometer (TOF-MS) (m/z (M+Na^+^), 617.2026; m/z (M+Na^+^_calc_), 617.2014).

### Biotin-2PCA and biotin-SS-2PCA modification of peptide libraries

Biotin-2PCA (**2**) and biotin-SS-2PCA (**3**) N-terminal modification reactions (50 μL total volume) contained 1 mg/mL (~1 mM) proteome-derived peptide library (25 μL of 2 mg/mL stock), 0.5 mM biotin-2PCA or biotin-SS-2PCA (2.5 μL of 10 mM stock in DMSO), and 50 mM sodium phosphate, pH 7.5 (6.3 μL of 400 mM stock). Reactions were initiated by addition of biotin-2PCA or biotin-SS-2PCA and were incubated at 37ºC for 2 h.

### Alkyne-2PCA modification of proteome-derived peptide libraries

Alkyne-2PCA (**4**) N-terminal modification reactions (50 μL total volume) contained 1 mg/mL (∼1 mM) proteome-derived peptide library (25 μL of 2 mg/mL stock), 0.5 mM alkyne-2PCA (2.5 μL of 10 mM stock in DMSO), and 50 mM sodium phosphate, pH 7.5 (6.3 μL of 400 mM stock). Reactions were initiated by addition of alkyne-2PCA and were incubated at 37ºC for 2 h.

### Click reactions for biotinylation of alkyne-2PCA-modified peptides

Click reactions (100 μL total volume) for biotinylation of alkyne-2PCA modified peptides contained 50 μL of alkyne-2PCA modification reaction mixture, 0.25 mM biotin-linker-azide (2.5 μL of a 10 mM DMSO stock; linkers were either DADPS (**8**), Dde (**7**), Diazo (**9**), disulfide (**6**), or no linker (**5**)), 0.4 mM CuSO4 and 0.8 mM BTTAA (3.2 μL of a solution containing 12.5 mM CuSO4 and 25 mM BTTAA), and 0.5 mg/mL sodium ascorbate (10 μL of a freshly made 5 mg/mL stock), and ddH2O to 100 μL. Components were added to the reaction in the following order: alkyne-modified peptide library, biotinylated azide, CuSO4/BTTAA, sodium ascorbate. Reactions were incubated for 1 h at 37ºC.

### Enrichment of biotinylated peptides

Biotinylated peptides were enriched on High-Capacity Neutravidin Agarose resin (ThermoFisher) (250 μL of 50% resin slurry per reaction). Click reactions were diluted with 4 volumes (400 μL) of 4 M guanidine hydrochloride and added to resin pre-equilibrated with 4 M guanidine hydrochloride. Samples were incubated for 30 min at room temperature on a rotisserie mixer. The resin slurry was transferred into a spin column (Pierce #69725) and attached to the QIAvac 24 plus vacuum manifold (Qiagen) for washing. The resin was washed ten times with 800 μL of 4 M guanidine hydrochloride and ten times with 800 μL of LC-MS grade water. Columns were removed from the vacuum manifold, the bottoms were capped, resin was resuspended in 225 μL of the appropriate elution buffer, and samples were transferred to 1.5 mL microcentrifuge tubes for elution. For biotin azide with no cleavable linker, peptides were eluted twice with 80% acetonitrile, 0.1% formic acid, first at 30ºC with shaking for 10 min, second at 72ºC with shaking for 10 min. The eluate was dried in a vacuum concentrator and redissolved in 1% TFA before desalting. For biotin-disulfide-azide, peptides were eluted with 5 mM TCEP for 1 h at room temperature. For biotin-DADPS-azide, peptides were eluted with 10% formic acid for 1 h at room temperature. For biotin-Diazo-azide, peptides were eluted with 25 mM dithionite in PBS for 1 h at room temperature. For biotin-Dde-azide, peptides were eluted with 2% aqueous hydrazine for 1 h at room temperature. Biotin-linker-azide samples were adjusted to pH <3 with TFA before desalting. Desalting was performed using SOLA HRP C18 columns (10 mg, Thermo Scientific). Columns were conditioned with 100% acetonitrile (500 μL) and equilibrated with 0.1% TFA (2 × 1 mL) before loading the sample. After sample loading, columns were washed with 0.1% TFA (2 × 1 mL), 0.1% TFA, 5% MeOH (1 mL), 0.1% formic acid, 2% acetonitrile (1 mL). Peptides were eluted with 50% acetonitrile, 0.1% formic acid (2 × 150 μL). The eluted peptides were dried in a vacuum concentrator and then redissolved in 0.1% formic acid prior to LC-MS/MS analysis.

### N-terminal reductive dimethylation of proteome-derived peptide libraries

N-terminal dimethylation reactions were carried out using a general method described previously^30^. N-terminal reductive dimethylation reactions (200 μL) contained 1 mg/mL peptide library, 30 mM formaldehyde (0.6 μL of a 10 M stock solution), 30 mM sodium cyanoborohydride (60 μL of a 100 mM stock solution), and 1% acetic acid (2 μL of glacial acetic acid). Reactions were allowed to proceed for 10 min at 37ºC and were quenched by loading onto a pre-conditioned and equilibrated SOLA HRP C18 cartridge by centrifugation at 100 × g for 1 min. The cartridge was then washed with 2 × 1 mL of 0.1% TFA and eluted with 300 μL 0.1% formic acid/50% acetonitrile. The solution was evaporated to dryness in a vacuum centrifuge and peptides were resuspended in water.

### Proteomic identification of cleavage sites with 2 PCA reagents (PICS2)

PICS2 reactions were performed with 100-400 μg of N-terminally blocked proteome-derived peptide library at a concentration of 1 mg/mL. Reaction buffer and protease amount varied. For trypsin, chymotrypsin, GluC, and LysargiNase PICS2 experiments, reactions were performed in 100 mM HEPES, pH 7.5 with a 1:100 (w/w) protease:peptide ratio. For Kex2 PICS2 experiments, reactions were performed in 100 mM Tris-HCl, pH 8.5 at a 1:20 protease:peptide ratio. Furin reactions were performed in 100 mM HEPES, pH 7.5, 1 mM CaCl2 at a 1:50 protease:peptide ratio. PCSK2 experiments were performed in 50 mM sodium acetate, pH 5.0, 100 mM NaCl, 1 mM CaCl2 at a 1:100 protease:peptide ratio. After 16 h, reactions were stopped by heating to 95ºC for 10 min. Reactions were allowed to cool and biotin-SS-2PCA was added to a final concentration of 0.5 mM from a 10 mM DMSO stock solution. The biotinylation reaction was allowed to proceed for 4-16 h at 37ºC. The reaction mixture was then diluted fivefold with 4 M guanidine hydrochloride and enriched on High-Capacity Neutravidin Agarose resin (1000 μL of 50% resin slurry per reaction) as described above. Biotin-SS-2PCA-modified peptides were eluted with 5 mM TCEP for 1-2 h at room temperature. Samples were desalted on SOLA HRP C18 cartridges prior to analysis by LC-MS/MS. PICS experiments performed for comparison were carried out as described previously58.

### Induction of apoptosis in Jurkat cells

Jurkat E6.1 (ATCC# TIB-152) cells were grown in RPMI-1640 media supplemented with 10% fetal bovine serum, 2 mM L-glutamine, and 1% penicillin-streptomycin at 37ºC under 5% CO2 atmosphere. One T225 flask of cells containing 250 mL of media with cells at a density of 106 cells per mL was used for each replicate. Cells were treated with either 50 μM etoposide (from a 50 mM DMSO stock solution) or an equal volume of DMSO for 8 h. Cells were harvested by centrifugation at 300 x g for 5 min and washed once with PBS. Jurkat were tested every six months for mycoplasma contamination using the LookOut Mycoplasma PCR Detection Kit (Sigma-Aldrich) according to the manufacturer’s instructions (**Fig. S25**).

### Alkyne-2PCA modification of cell lysates

Each cell pellet was resuspended in 800 μL hot lysis buffer (100 mM Tris-HCl, pH 8.5, 6 M guanidine hydrochloride, 5 mM TCEP, 10 mM chloroacetamide) that had been preheated to 95ºC and the suspension was heated at 95ºC for 10 min. Lysis was completed using 10 cycles of probe ultrasonication (20% amplitude; 5 s on / 5 s off). Insoluble material was removed by centrifugation at 20,000 × g at 4ºC for 10 min. After cooling, alkyne-2PCA (100 mM stock solution in DMSO) was added the supernatants to a final concentration of 1 mM and the lysate was incubated overnight at 37ºC. Protein was then precipitated by addition of 15% (w/v) trichloroacetic acid and incubation for 16 h at -20ºC. Precipitated protein was pelleted by centrifugation at 20,000 × g for 5 min and the supernatant was removed. Pellets were washed two times with 200 μL ice-cold acetone, centrifuged at 20,000 × g for 5 min, and air dried. Pellets were redissolved in 800 μL 20 mM NaOH. Cu(I)-catalyzed azide-alkyne cycloaddition (CuAAC) was the performed by mixing 257.2 μL of the protein solution with 800 μL 1 M HEPES (200 mM final concentration), 10 μL 10 mM biotin-SS-azide (0.25 mM final concentration), 12.8 μL of a pre-mixed solution containing 12.5 mM CuSO4 and 25 mM BTTAA (0.4 mM CuSO4 and 0.8 mM BTTAA final concentration), and 40 μL of 5 mg/mL freshly prepared sodium ascorbate. Click reactions were incubated for 1 h at 37ºC. After 1 h, the reaction mixture was diluted with three volumes (1.2 mL) of 4 M GdnHCl and added to 250 μL of High-Capactiy Neutravidin Agarose resin that had been pre-equilibrated with 4 M GdnHCl. Samples were incubated for 30 min at room temperature on a rotisserie mixer. The resin slurry was transferred into a spin column (Pierce #69725) and attached to the QIAvac 24 plus vacuum manifold (Qiagen) for washing. The resin was washed ten times with 800 μL of 4 M guanidine hydrochloride and ten times with 800 μL of 20 mM HEPES, pH 7.5. Columns were removed from the vacuum manifold, the bottoms were capped, resin was resuspended in 500 μL of 20 mM HEPES, pH 7.5, and samples were transferred to 1.5 mL microcentrifuge tubes. Trypsin (20 μg) was addition and on-bead digestion was allowed to proceed overnight at room temperature on a rotisserie mixer. Following digestion, the slurry was transferred into a spin column and attached to the for washing. The resin was washed ten times with 800 μL of 4 M guanidine hydrochloride and ten times with 800 μL of 20 mM HEPES, pH 7.5. Columns were removed from the vacuum manifold, the bottoms were capped, resin was resuspended in 500 μL of 20 mM HEPES, pH 7.5, and samples were transferred to 1.5 mL microcentrifuge tubes. TCEP was added to a final concentration of 5 mM and samples were incubated for 2 h at room temperature on a rotisserie mixer. After TCEP reduction, the resin slurry was transferred to a spin column and eluted N-terminal peptides were collected by centrifugation at 500 × g for 2 min. Samples were acidified by addition of TFA to 1% and desalted using SOLA HRP C18 desalting cartridges. Desalted peptides were dried in a vacuum centrifuge.

### LC-MS/MS data collection

Dried peptide samples were redissolved in 0.1% formic acid prior to LC-MS/MS analysis. Peptide concentration was estimated by absorbance at 280 nm assuming that 1 abosrbance unit = 1 mg/mL. For each experiment, 500 ng of peptides were analyzed with an Orbitrap Exploris 480 hybrid quadrupole-Orbitrap mass spectrometer coupled to an UltiMate 3000 RSLCnano liquid chromatography system (ThermoFisher Scientific). Peptides were loaded onto an Acclaim PepMap RSLC column (75 μm × 15 cm, 2 μm particle size, 100 Å pore size, ThermoFisher Scientific) over 15 min in 97% mobile phase A (0.1% formic acid) and 3% mobile phase B (0.1% formic acid, 80% acetonitrile) at 0.5 μL/min. Peptides were eluted at 0.3 μL/min using a linear gradient from 3% mobile phase B to 50% mobile phase B over 120 min. Peptides were electrosprayed through a nanospray emitter tip connected to the column by applying 2000 V through the ion source’s DirectJunction adapter. Full MS scans were performed at a resolution of 60,000 at 200 m/z over a range of 300-1,200 m/z with an AGC target of 300% and the maximum injection time set to ‘auto’. The top 20 most abundant precursors with a charge state of 2-6 were selected for MS/MS analysis with an isolation window of 1.4 m/z and a precursor intensity threshold of 5 × 103. A dynamic exclusion window of 20 s with a precursor mass tolerance of ± 10 ppm was used. MS/MS scans were performed using HCD fragmentation using a normalized collision energy of 30% and a resolution of 15,000 with a fixed first mass of 110 m/z. The AGC target was set to ‘standard’ with a maximum injection time of 22 ms.

### Mass spectrometry data analysis

Open and offset searches were performed in FragPipe 17.1. For open searches, the precursor mass tolerance was set to -150-500 Da. The initial fragment mass tolerance was set to 0.02Da. Calibration and Optimization was set to ‘Mass calibration, parameter optimization’. Isotope error was set to 0. Cleavage was set to ‘Enzymatic’ with clip N term enabled and enzyme name was set to trypsin, chymotrypsin, or gluc as appropriate. Up to two missed cleavages were allowed. Peptide length was set to 7-50 and peptide mass range was set to 500-5,000 Da. No variable modifications or fixed modifications were specified. Crystal-C was enabled and PeptideProphet was run with the following settings: ‘--nonparam --expectscore --decoyprobs --masswidth 1000.0 --clevel -2’. PTMProphet was disabled. The numbers of peptide-spectrum matches (PSMs) reported in the global.modsummary.tsv file were then plotted against the reported mass shift. Results can be found in **Dataset S2**.

For offset searched in FragPipe, the precursor mass tolerance was set to 10 ppm and the fragment mass tolerance was set to 0.02 Da. Calibration and Optimization was set to ‘Mass calibration, parameter optimization’. Isotope error was set to 0/1/2/3. Cleavage was set to ‘Enzymatic’ with clip N term enabled and enzyme name was set to trypsin, chymotrypsin, or gluc as appropriate. Up to two missed cleavages were allowed. Peptide length was set to 7-50 and peptide mass range was set to 500-5,000 Da. Oxidation at Met (+15.9949) and acetylation at protein N termini (+42.0106) were searched as variable modification and Cys carbamidomethylation (+57.02146) was searched as a fixed modification. Mass offsets of 0 and 89.0265 were included with Restrict delta mass set to all. Crystal-C was disabled. PeptideProphet was run with the following settings” ‘--decoyprobs --ppm --accmass --nonparam --expectscore – database databasepath’, where database path points to the fasta database that was searched. Results can be found in **Dataset S2**.

Thermo RAW files were searched against either the human or *E*. coli SwissProt database using the SEQUEST algorithm in Proteome Discoverer 2.4 (Thermo). The precursor mass tolerance was set at 10 ppm and the fragment mass tolerance was set at 0.02 Da. Search parameters included carbamidomethylation at Cys (+57.021 Da) as a constant modification and the following dynamic modifications: oxidation at Met (+15.995 Da), acetylation at protein N termini (+42.011 Da), Met loss at protein N termini (−131.040 Da), Met loss+acetylation at protein N termini (−89.030 Da). Additional dynamic modifications at the peptide N terminus were included as appropriate for the experiment. Modification masses are listed in **Table S2**. Up to two missed cleavages were allowed. For PICS and CHOPPER data analysis, cleavage specificity was set to semi (C-term), requiring that the C terminus of each peptide have the specificity of the digest protease but allowing the N terminus to vary. The Percolator node of Proteome Discoverer was used for PSM validation at a false discovery rate of 1%. Raw datafiles, peak lists and results have been deposited in the ProteomeXchange repository under the accession numbers listed in **Table S3**. Results are also provided in Microsoft Excel format as **Datasets S1-S27** as described in **Table S3**.

### Specificity data analysis

Enrichment scores for 2PCA specificity were calculated using custom Python scripts available in the GitHub repository. These scores correspond to the standard score (z score) comparing frequencies of amino acids in the total population of peptides to frequencies of amino acids in the 2PCA-modified peptide population and were calculated using the equation

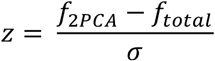

where f_2PCA_ is the frequency of the amino acid among the 2PCA-modified population, f_total_ is the frequency of the amino acid in the total population of peptides, and σ is the population standard deviation.

### PICS2 data analysis

The frequencies of each amino acid at each position in PICS samples were calculated using a custom Python script that is available in the GitHub repository. This script filters Proteome Discoverer search results for peptides that are modified with cleaved biotin-SS-2PCA and whose cleavage sites do not match the specificity of the protease used for library generation. It then retrieves the protein sequence from a SwissProt xml file containing proteome sequences for either human or *E. coli* and uses this information to infer the nonprime side sequence of the measured peptide (i.e., the sequence N-terminal to the cleavage site). The script then counts the number of times each amino acid is observed in each position and calculates a frequency by dividing by the total number of observations. IceLogos were constructed using iceLogo1.259 with either the human or *E. coli* SwissProt database as a reference.

### CHOPPER data analysis

CHOPPER data analysis was performed using a custom Python script that is available in the GitHub repository. This script filters Proteome Discoverer search results for peptides that are modified with alkyne-2PCA clicked to cleaved biotin-2PCA. It then retrieves the protein sequence from a SwissProt60 xml file containing all human proteome protein sequences to infer the nonprime side residues of the cleavage site. Cleavage sites were plotted on domain boundaries using a custom Python script that retrieves domain boundaries from the SwissProt xml file in combination with matplotlib. STRING analysis was performed using Cytoscape 3.9.161. with a confidence score cutoff of 0.99. All proteins known to be cleaved with P1 = D specificity upon etoposide treatment of Jurkat cells were used in the analysis. **Figure 5G** shows only clusters that contain at least one cleavage identified in the CHOPPER dataset. **Fig. S21** shows the full network of STRING interactions.

## Code availability

Python scripts used for data analysis are available in the GitHub repository (http://dx.doi.org/10.5281/zenodo.3678926) under GNU General Public License v3.0.

## Data availability

All data generated for this study are available in the main text or Supporting Information. In addition, raw MS data and search results have been deposited in ProteomeXchange under the accession numbers listed in **Table S3**.

