## Supplementary Figures and Tables for "An expanded 2-pyridinecarboxaldehyde (2PCA)-based chemoproteomics toolbox for probing protease specificity"

**Affiliation**

Supplementary Information contains Figures S1-S6 and Tables S1-S3. Supplementary Datasets 1-27 have been included as separate Microsoft Excel files.

**Figure S1. IceLogos for *E. coli* proteome-derived peptide libraries.** (A) IceLogo for *E. coli* proteome-derived peptide library generated with chymotrypsin. (B) IceLogo for *E. coli* proteome-derived peptide library generated with GluC. (C) IceLogo for *E. coli* proteome-derived peptide library generated with trypsin. IceLogos were generated using IceLogo 1.2 using the first six amino acids of each peptide identified in the library and the C-terminal residue as the experimental set and the pre-compiled SwissProt composition for E. coli K12 as the reference set.

A

*E. coli* chymotryptic peptide library (n = 13,036 peptides)

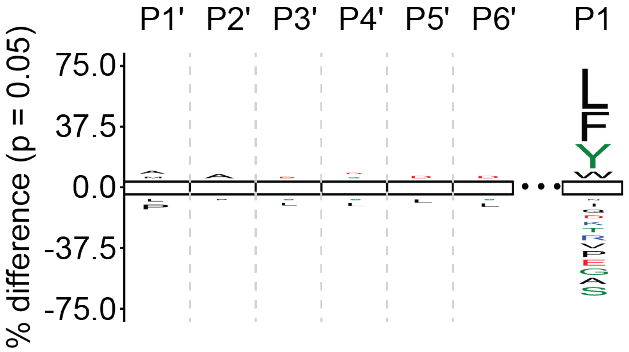


B

*E. coli* GluC peptide library (n = 8,252 peptides)

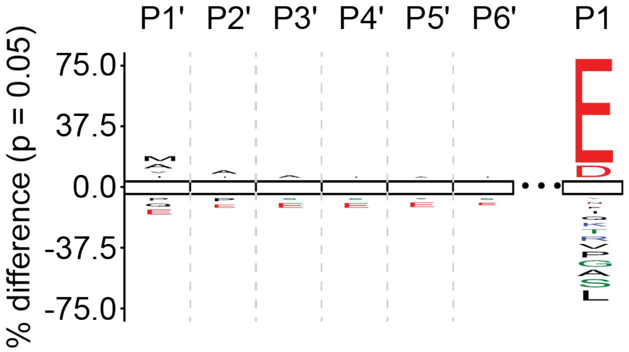


C

*E. coli* trypsin peptide library (n = 19,601 peptides)

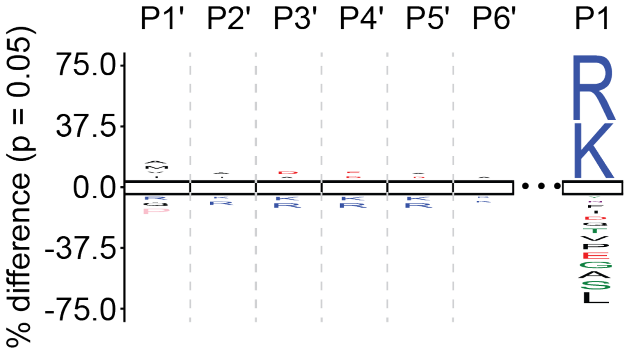

**Figure S2. Individual 2PCA-modified *E. coli* proteome-derived peptide libraries.** (A) 2PCA-modified *E. coli* trypsin library. (B) 2PCA-modified *E. coli* GluC library. (C) 2PCA-modified *E. coli* chymotrypsin library.

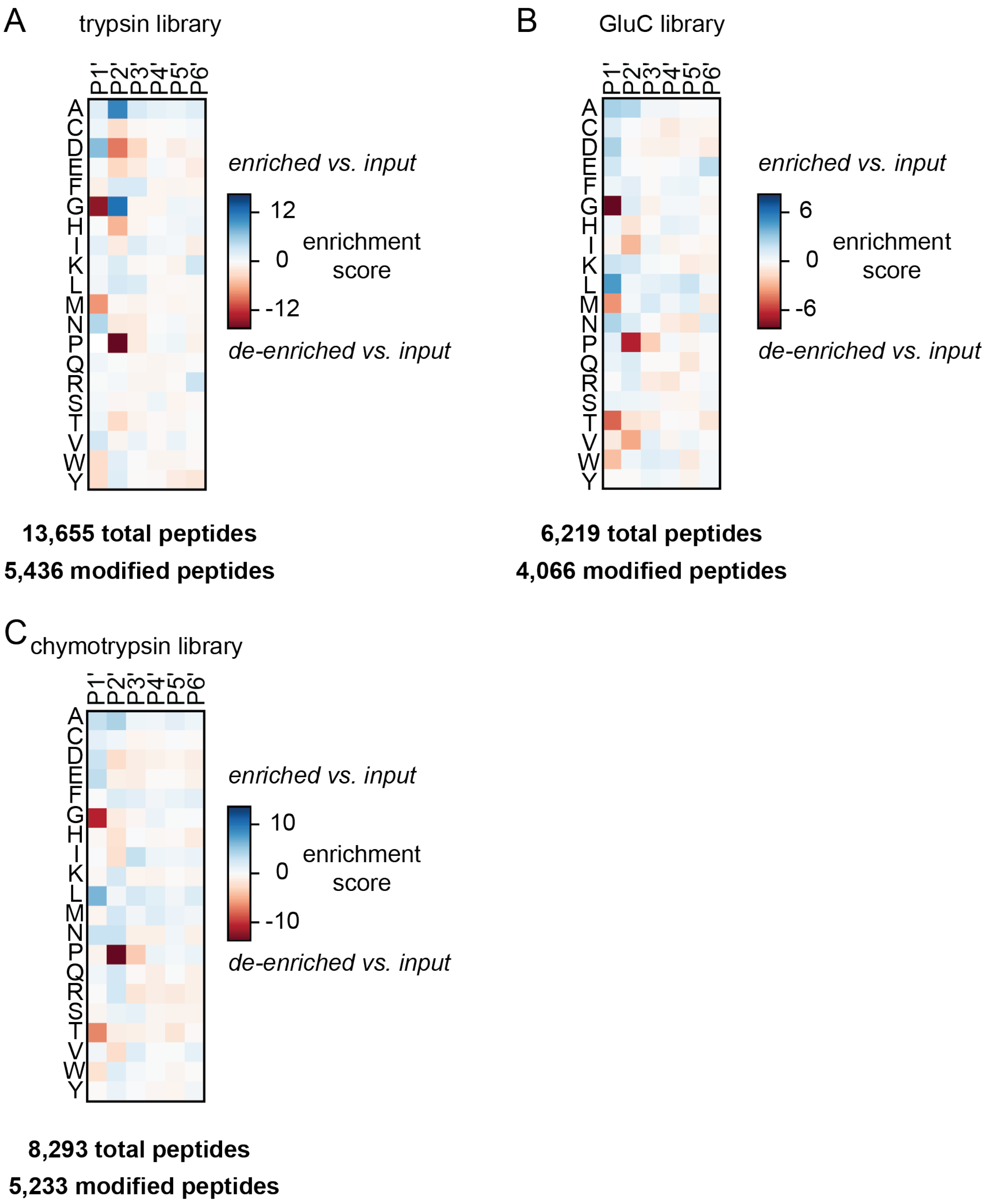

**Figure S3. MS/MS spectra for putative P2' proline peptides.** (A) MS/MS spectrum and predicted fragment ion m/z for the peptide 2PCA-HPELTDMVIFR. (B) MS/MS spectrum and predicted fragment ion m/z for the peptide 2PCA- 2PCA-SPVEATLR. Matched y-ions are colored in blue and matched b-ions are colored in red.

A

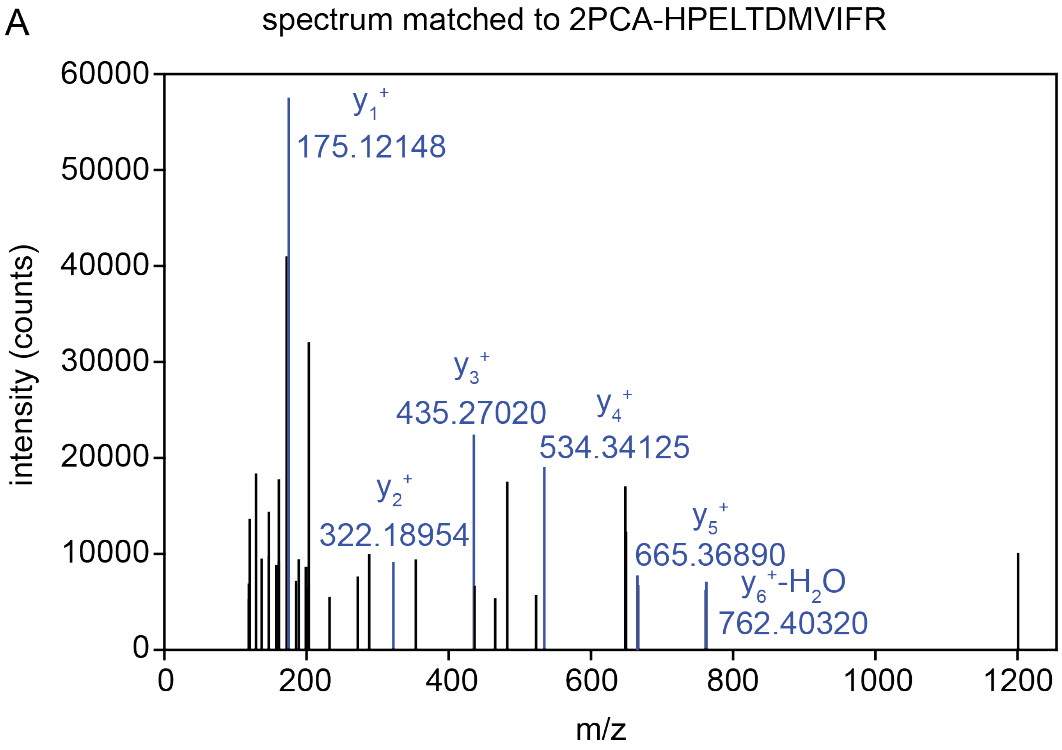


| sequence | b-ion # | b | y | y-ion # |
|---|---|---|---|---|
| H(2PCA) | 1 | 227.0928 | --- | 11 |
| P | 2 | 324.1456 | 1220.6344 | 10 |
| E | 3 | 453.1881 | 1123.5816 | 9 |
| L | 4 | 566.2722 | 994.5390 | 8 |
| T | 5 | 667.3199 | 881.4550 | 7 |
| D | 6 | 782.3468 | 780.4073* | 6 |
| M | 7 | 913.3873 | 665.3803 | 5 |
| V | 8 | 1012.4557 | 534.3398 | 4 |
| I | 9 | 1125.5398 | 435.2714 | 3 |
| F | 10 | 1272.6082 | 322.1874 | 2 |
| R | 11 | 1428.7094 | 175.1190 | 1 |

*match to $y_6^+$-$H_2O$: 762.3967

**Figure S3 (cont'd).**

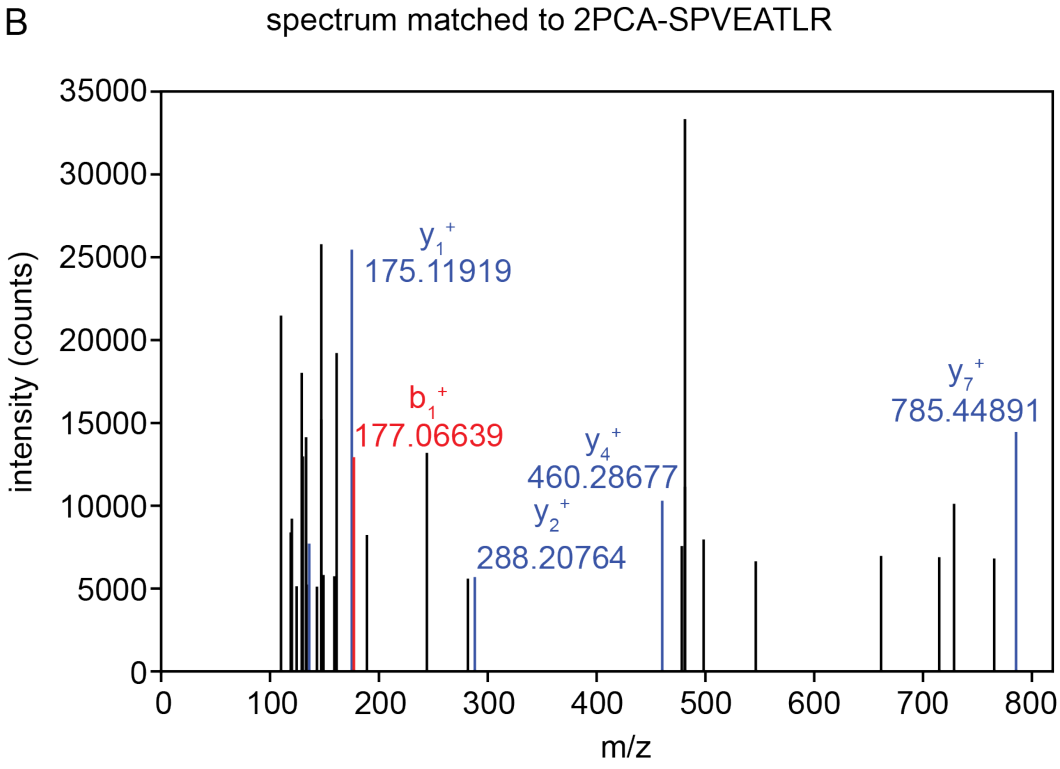


| sequence | b-ion # | b | y | y-ion # |
|---|---|---|---|---|
| S(2PCA) | 1 | 177.0660 | 961.5103 | 8 |
| P | 2 | 274.1187 | 785.4516 | 7 |
| V | 3 | 373.1871 | 688.3988 | 6 |
| E | 4 | 502.2297 | 589.3304 | 5 |
| A | 5 | 573.2668 | 460.2878 | 4 |
| T | 6 | 674.3145 | 389.2507 | 3 |
| L | 7 | 787.3985 | 288.2030 | 2 |
| R | 8 | 943.4997 | 175.1190 | 1 |

**Figure S4. Pairwise analysis of positional amino acid identity in 2PCA-modified peptides.** Enrichment or de-enrichment of pairwise combinations of amino acids in 2PCA-modified peptides was examined and enrichment scores were calculated as described in Methods.

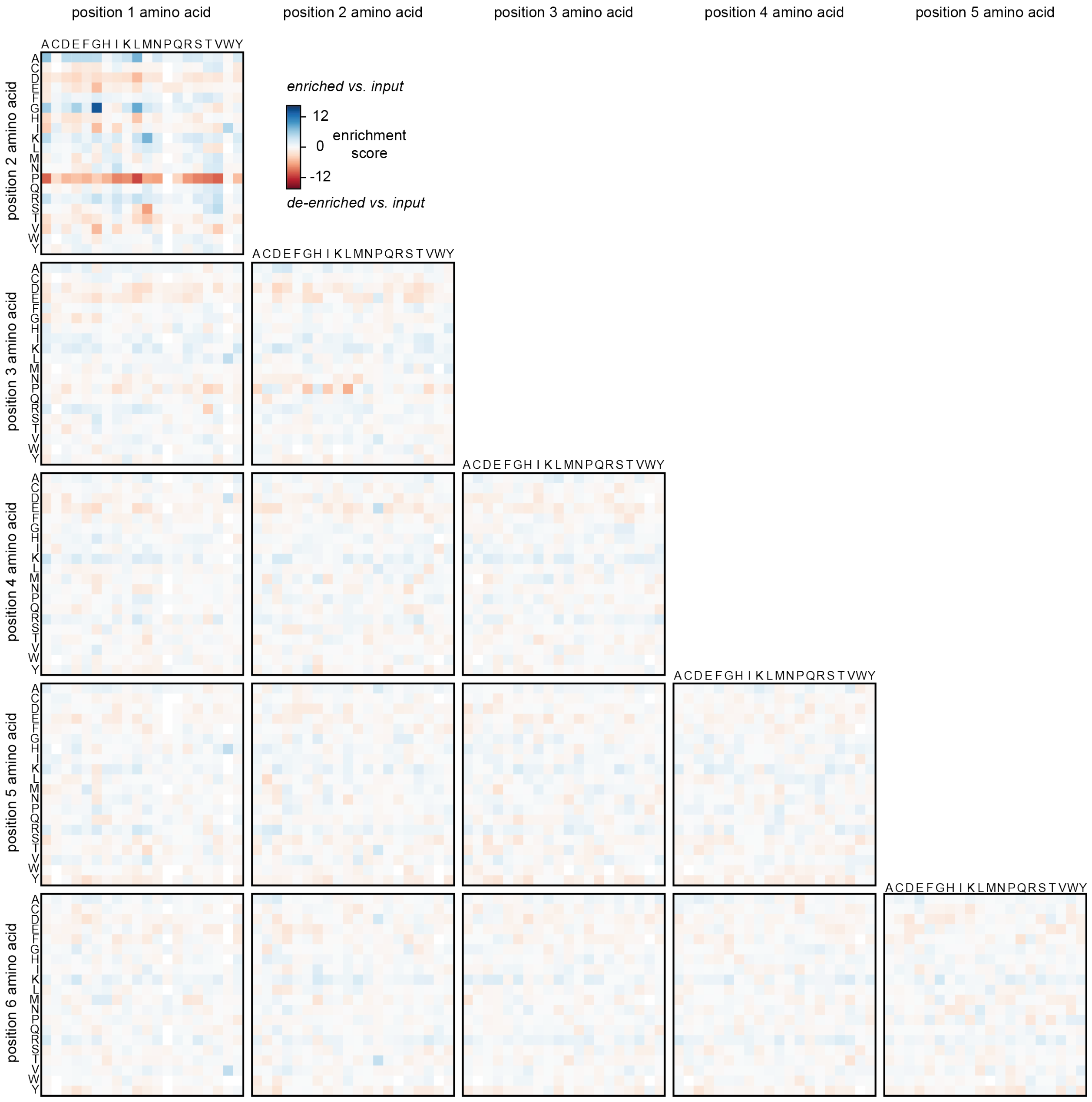

**Figure S5. Concentration dependence of 2PCA modification efficiency for individual peptide libraries.**

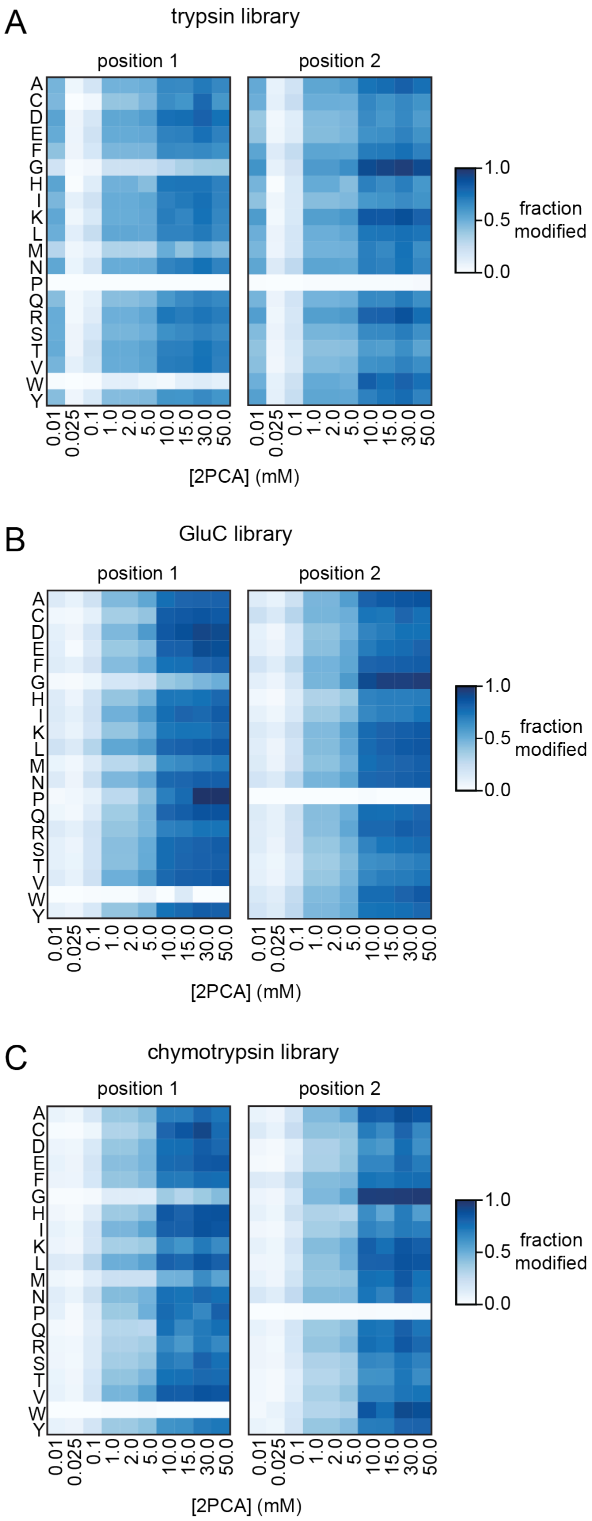

**Figure S6. Concentration dependence of 2PCA specificity.**

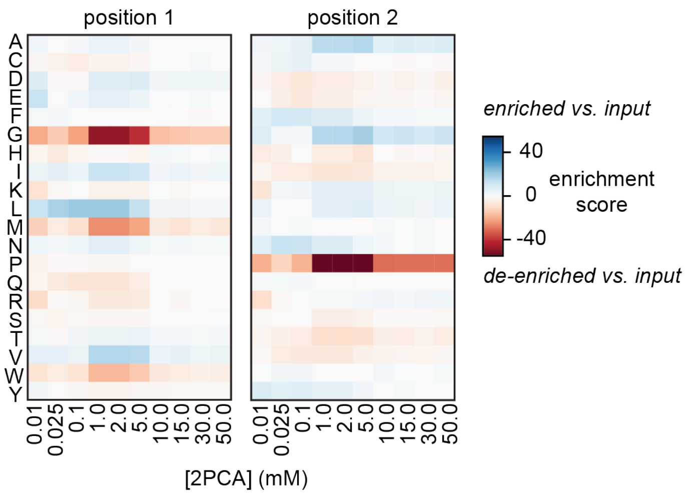

**Figure S7. Alkyne-2PCA modification of proteome-derived peptide libraries.** (A) Efficiency of alkyne-2PCA proteome-derived peptide library modification. (B) Residue specificity of alkyne-2PCA proteome-derived peptide modification.

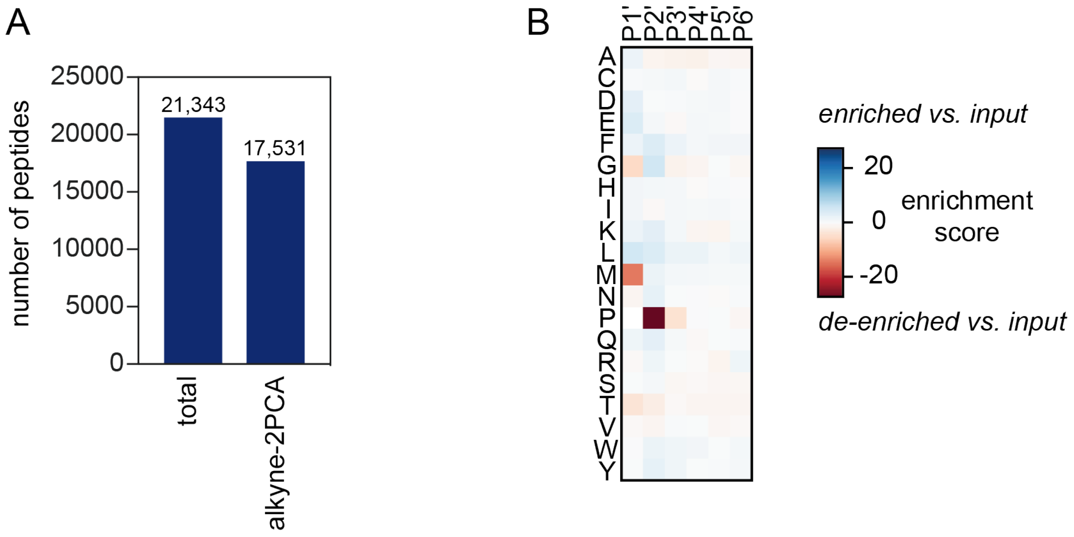

**Figure S8. Efficiency of Cu(I)-catalyzed azide-alkyne cycloaddition (CuAAC) on alkyne-2PCA proteome-derived peptide libraries using biotin azide.**

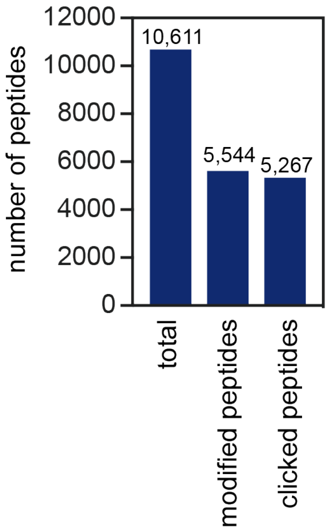

**Figure S9. IceLogos for human proteome-derived peptide libraries.** (A) IceLogo for human proteome-derived peptide library generated with trypsin. (B) IceLogo for human proteome-derived peptide library generated with GluC. (C) IceLogo for human proteome-derived peptide library generated with chymotrypsin. IceLogos were generated using IceLogo 1.2 using the first six amino acids of each peptide identified in the library and the C-terminal residue as the experimental set and the pre-compiled SwissProt composition for human as the reference set.

A *Human* trypsin peptide library (n = 33,437 peptides)

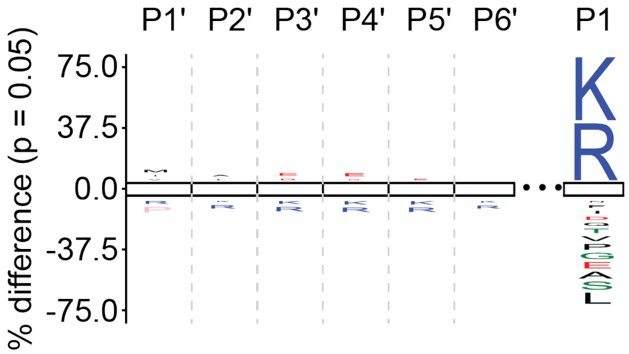


B *Human* GluC peptide library (n = 11,330 peptides)

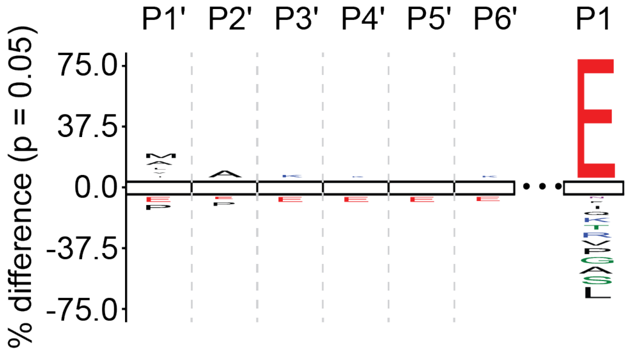


C *Human* chymotryptic peptide library (n = 7,288 peptides)

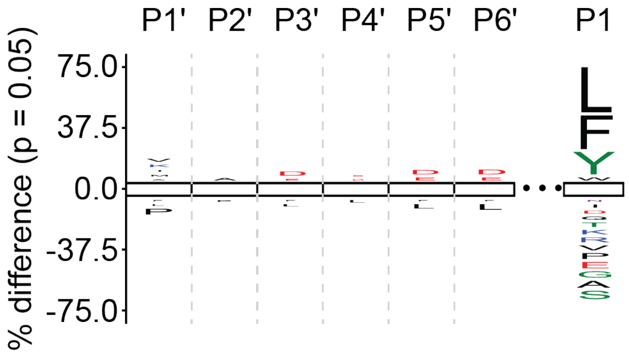

**Figure S10. Efficiency of 2PCA N-terminal blocking of proteome-derived peptide libraries.** Triangles represent trypsin libraries, circles represent chymotrypsin libraries, and squares represent GluC libraries.

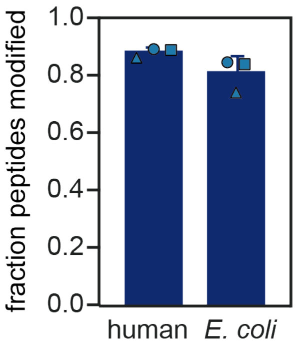

**Figure S11. Trypsin PICS and PICS2 analysis on individual proteome-derived peptide libraries.** (A) PICS with *E. coli*-derived peptide libraries dimethylated on N termini and lysine residues. (B) PICS2 with *E. coli*-derived peptide libraries N-terminally modified with 2PCA. (C) PICS with human-derived peptide libraries N-terminally modified with 2PCA.

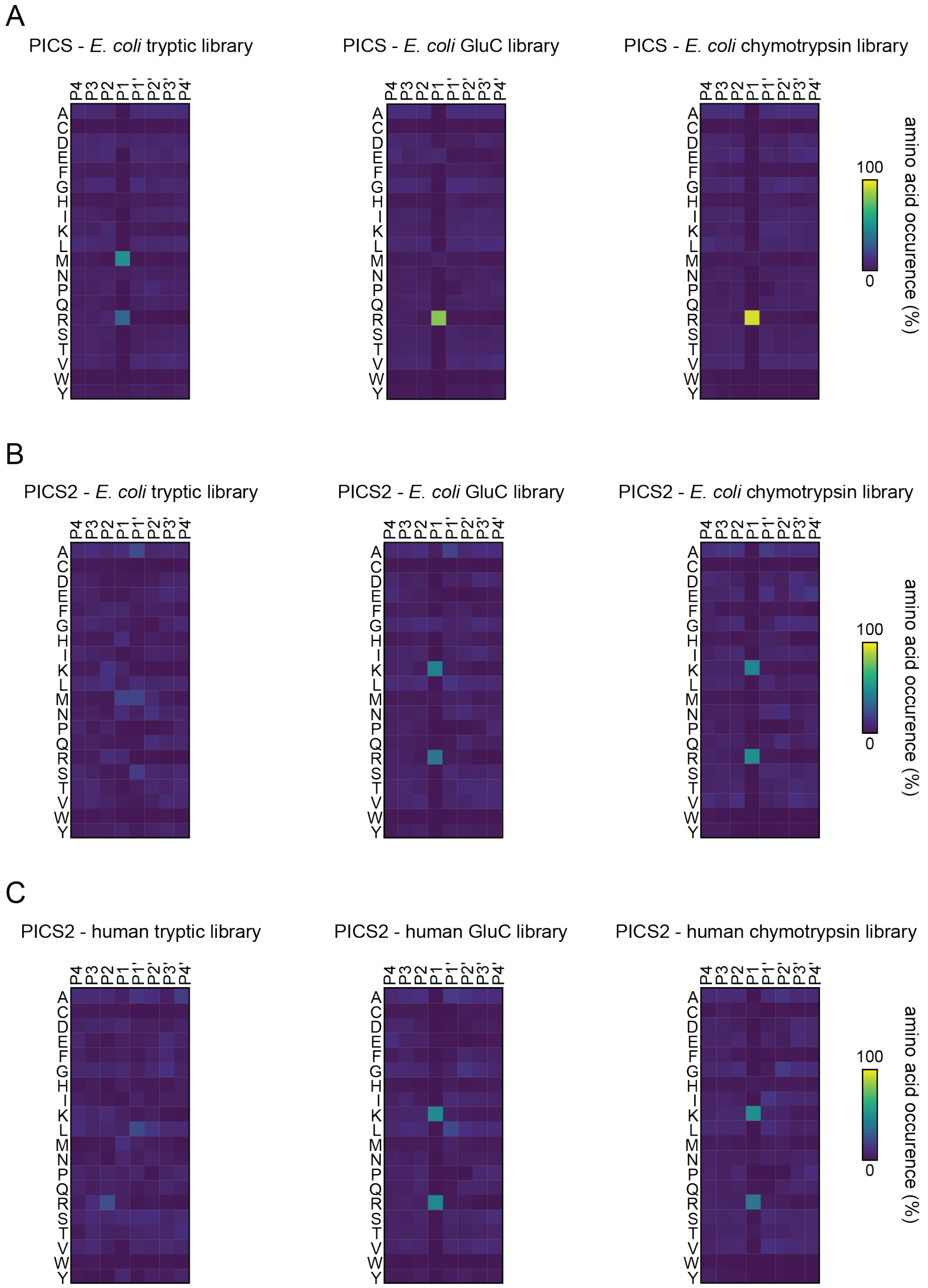

**Figure S12. LysargiNase PICS2 analysis on individual proteome-derived peptide libraries.**

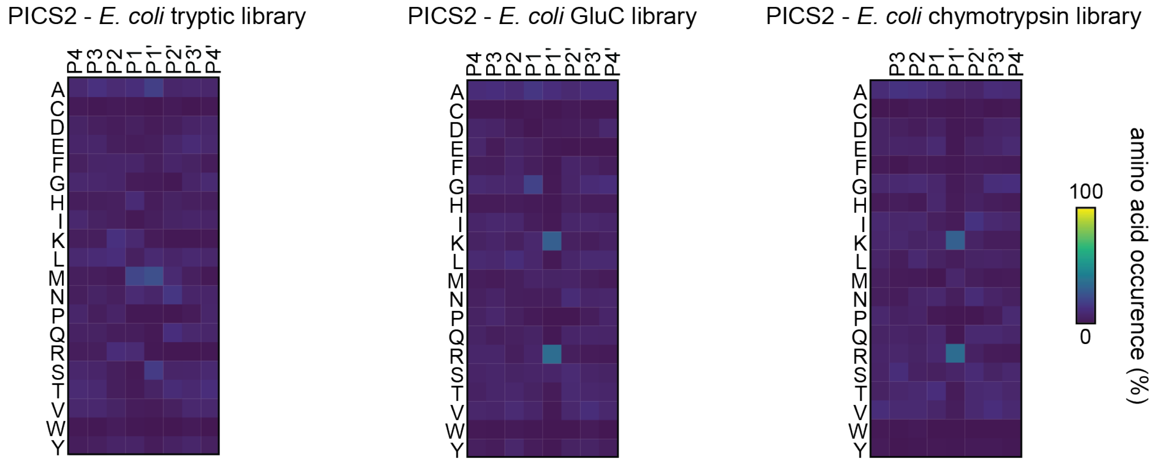

**Figure S13. GluC PICS2 analysis.** PICS2 was performed using an N-terminally 2PCA modified proteome-derived peptide library generated with trypsin from an *E. coli* protein extract.

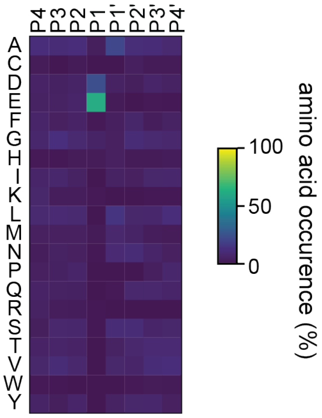

**Figure S14. Chymotrypsin PICS2 analysis.** PICS2 was performed using an N-terminally 2PCA modified proteome-derived peptide library generated with trypsin from an *E. coli* protein extract.

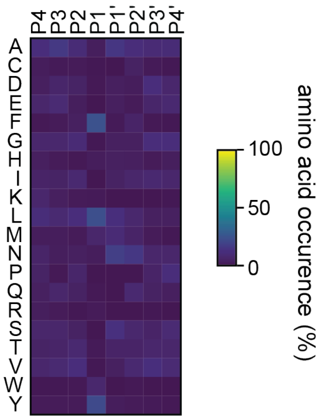

**Figure S15. Efficiency and specificity of reductive dimethylation under acidic conditions for N-terminal blocking.**

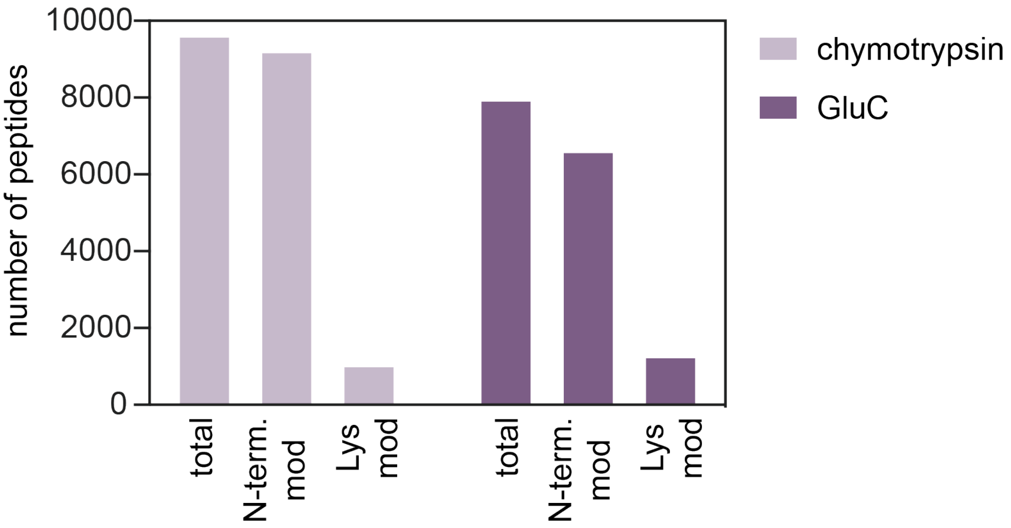

**Figure S16. Kex2 PICS2 analysis on individual proteome-derived peptide libraries.** Analysis was performed with *E. coli* proteome-derived peptide libraries N-terminally blocked by dimethylation.

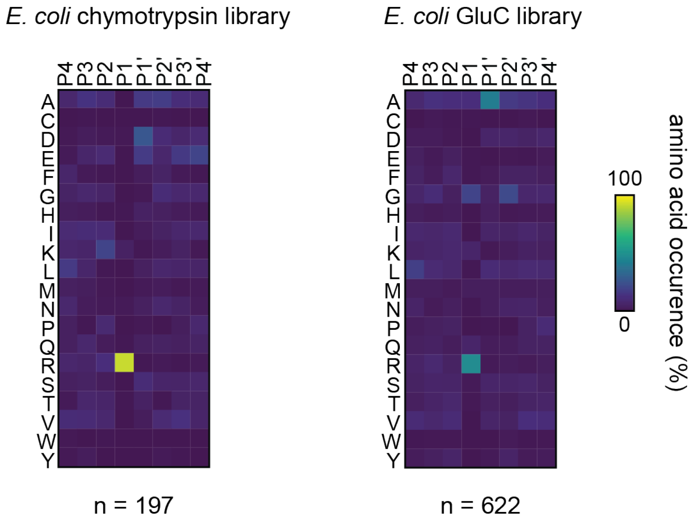

**Figure S17. Previously identified substrates of Kex2.** Kex2 was previously known to cleave pro-α-factor and K1 killer toxin. Whether Kex2 directly cleaves the PR site in killer toxin was previously unknown.

P01149 - Mating factor alpha-1

```
        10         20         30         40         50         60
MRFPSIFTAV LFAASSALAA PVNTTTEDET AQIPAEAVIG YLDLEGDFDV AVLPFSNSTN

        70         80         90        100        110        120
NGLLFINTTI ASIAAKEEGV SLDKREAEAW HWLQLKPGQP MYKREAEAEA WHWLQLKPGQ

       130        140        150        160   165
PMYKREADAE AWHWLQLKPG QPMYKREADA EAWHWLQLKP GQPMY
```

P01546 - M1-1 protoxin (K1 killer toxin)

```
        10         20         30         40         50         60
MTKPTQVLVR SVSILFFITL LHLVVALNDV AGPAETAPVS LLPREAPWYD KIWEVKDWLL

        70         80         90        100        110        120
QRATDGNWGK SITWGSFVAS DAGVVIFGIN VCKNCVGERK DDISTDCGKQ TLALLVSIFV

       130        140        150        160        170        180
AVTSGHHLIW GGNRPVSQSD PNGATVARRD ISTVADGDIP LDFSALNDIL NEHGISILPA

       190        200        210        220        230        240
NASQYVKRSD TAEHTTSFVV TNNYTSLHTD LIHHGNGTYT TFTTPHIPAV AKRYVYPMCE

       250        260        270        280        290        300
HGIKASYCMA LNDAMVSANG NLYGLAEKLF SEDEGQWETN YYKLYWSTGQ WIMSMKFIEE

       310    316
SIDNANNDFE GCDTGH
```

**Figure S18. Furin PICS2 analysis on individual proteome-derived peptide libraries.** Analysis was performed using human proteome-derived peptide libraries N-terminally blocked with 2PCA.

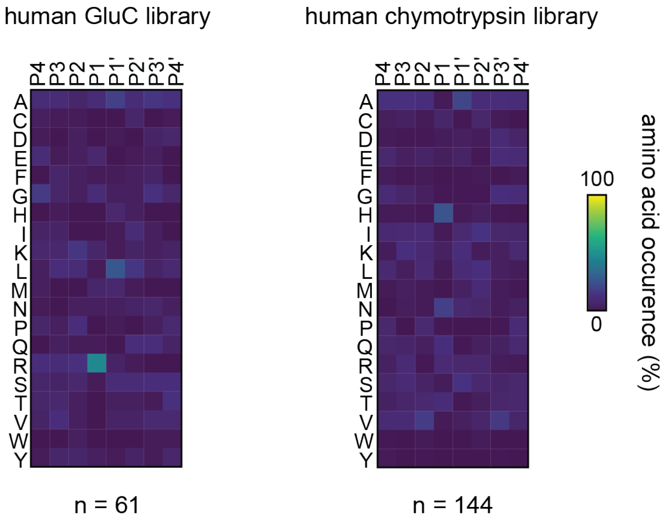

**Figure S19. PCSK2 PICS2 analysis on individual proteome-derived peptide libraries.** Analysis was performed using human proteome-derived peptide libraries N-terminally blocked by dimethylation.

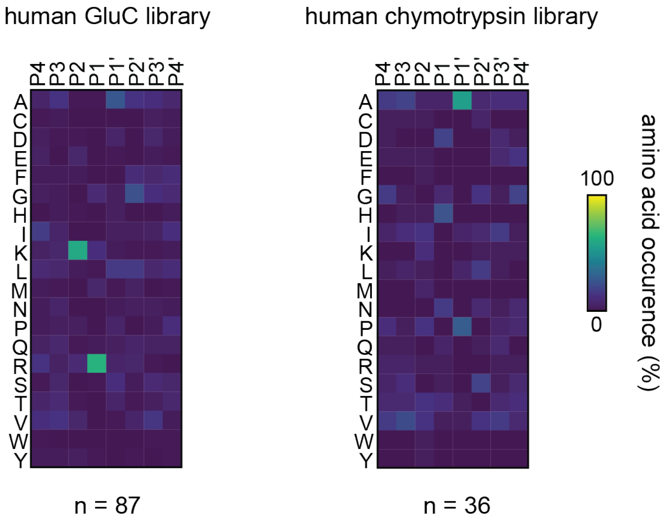

**Figure S20. Timecourse for etoposide treatment of Jurkat cells with etoposide or DMSO vehicle.** Images were collected at 0, 2, and 8 h after addition of etoposide or DMSO. At 8 h, etoposide-treated cells show membrane blebs characteristic of cells undergoing apoptosis while DMSO-treated cells have a much lower extent of blebbing.

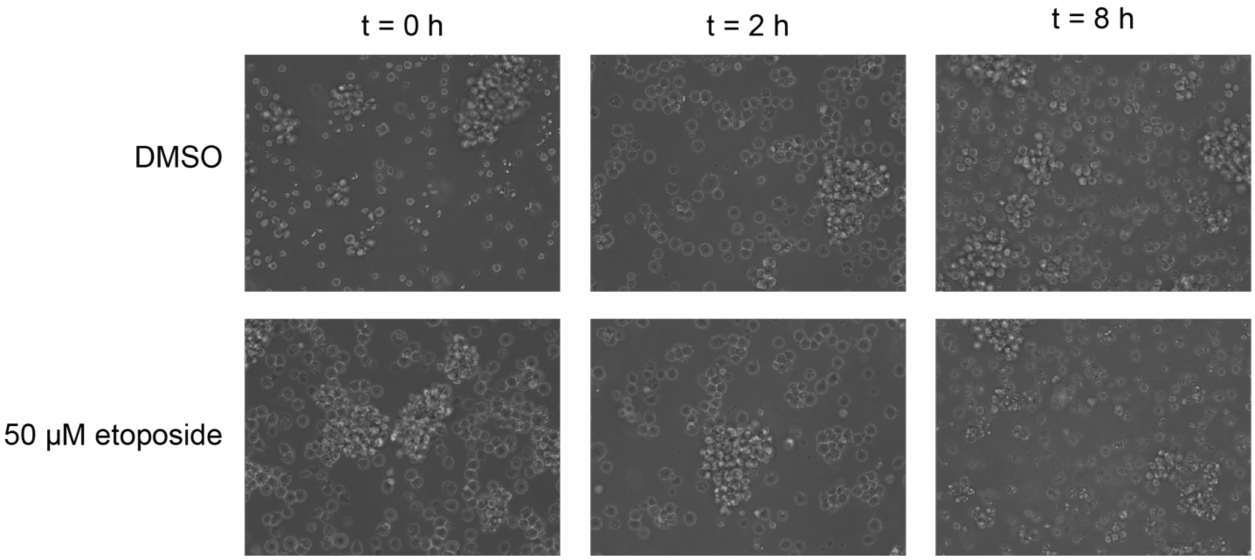

**Figure S21. STRING analysis of proteins that undergo caspase cleavage during apoptosis.** Cleaved proteins that were previously observed (grey) or newly identified by CHOPPER (magenta) were analyzed. Only proteins that have at least one interaction are shown.

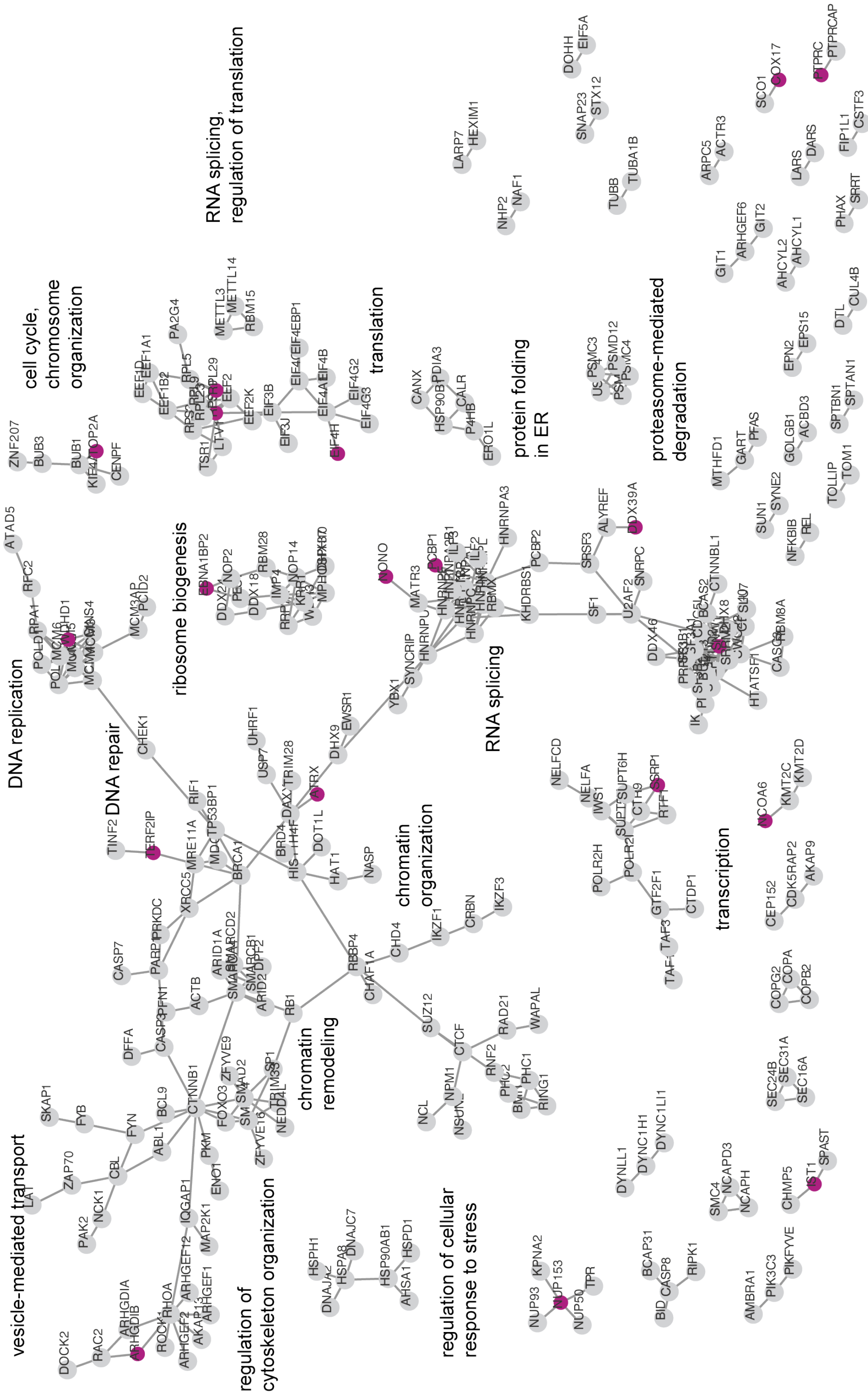

**Figure S22. Caspase cleavage sites identified with CHOPPER plotted with protein domain boundaries.** Domain boundaries were extracted from the Uniprot Knowledge base. Cleavage sites that were newly identified by CHOPPER are shown as magenta arrows and previously identified cleavage sites are shown as black arrows. Proteins are labeled at the top right with their Uniprot ID.

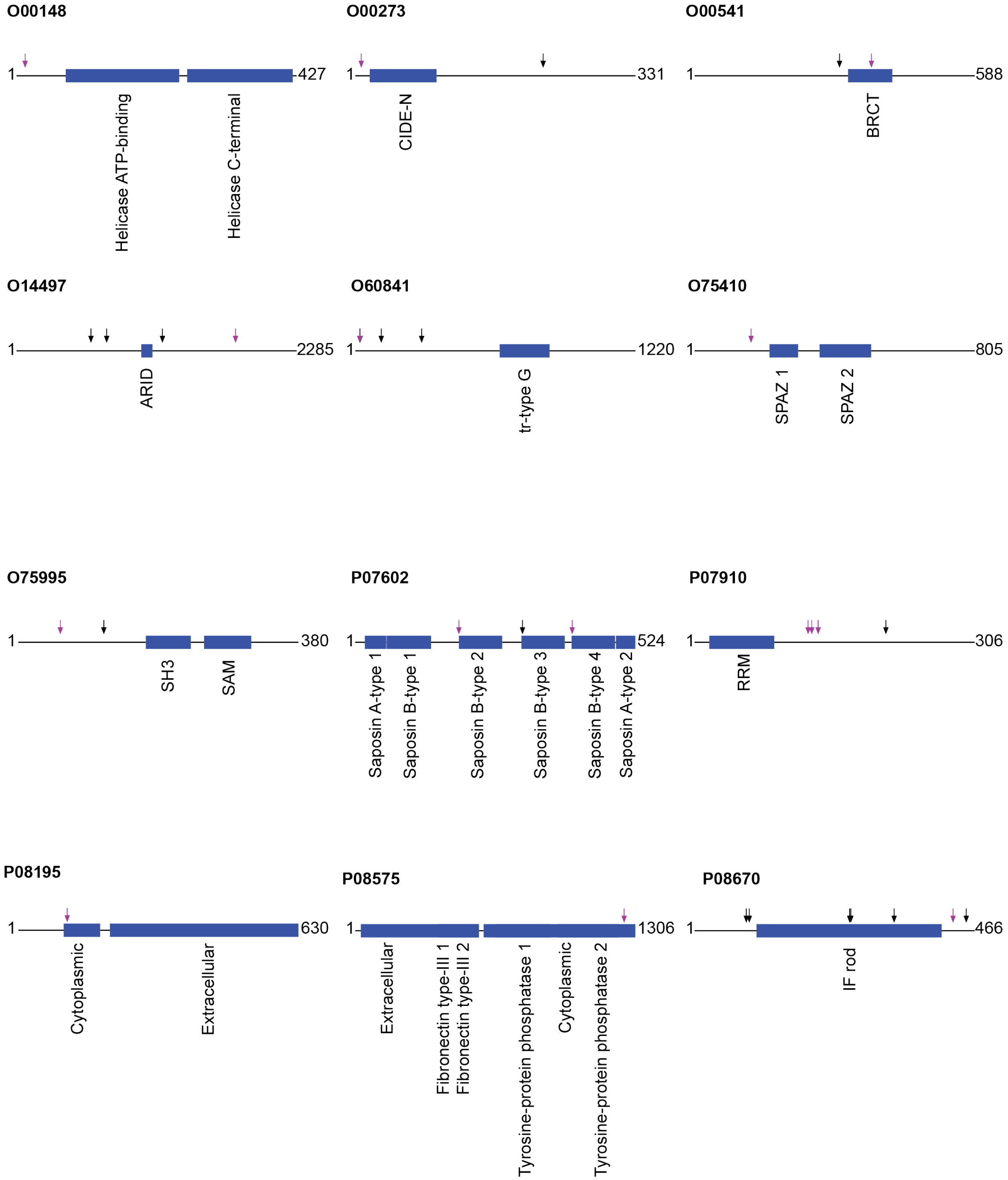

**Figure S22 (cont'd).**

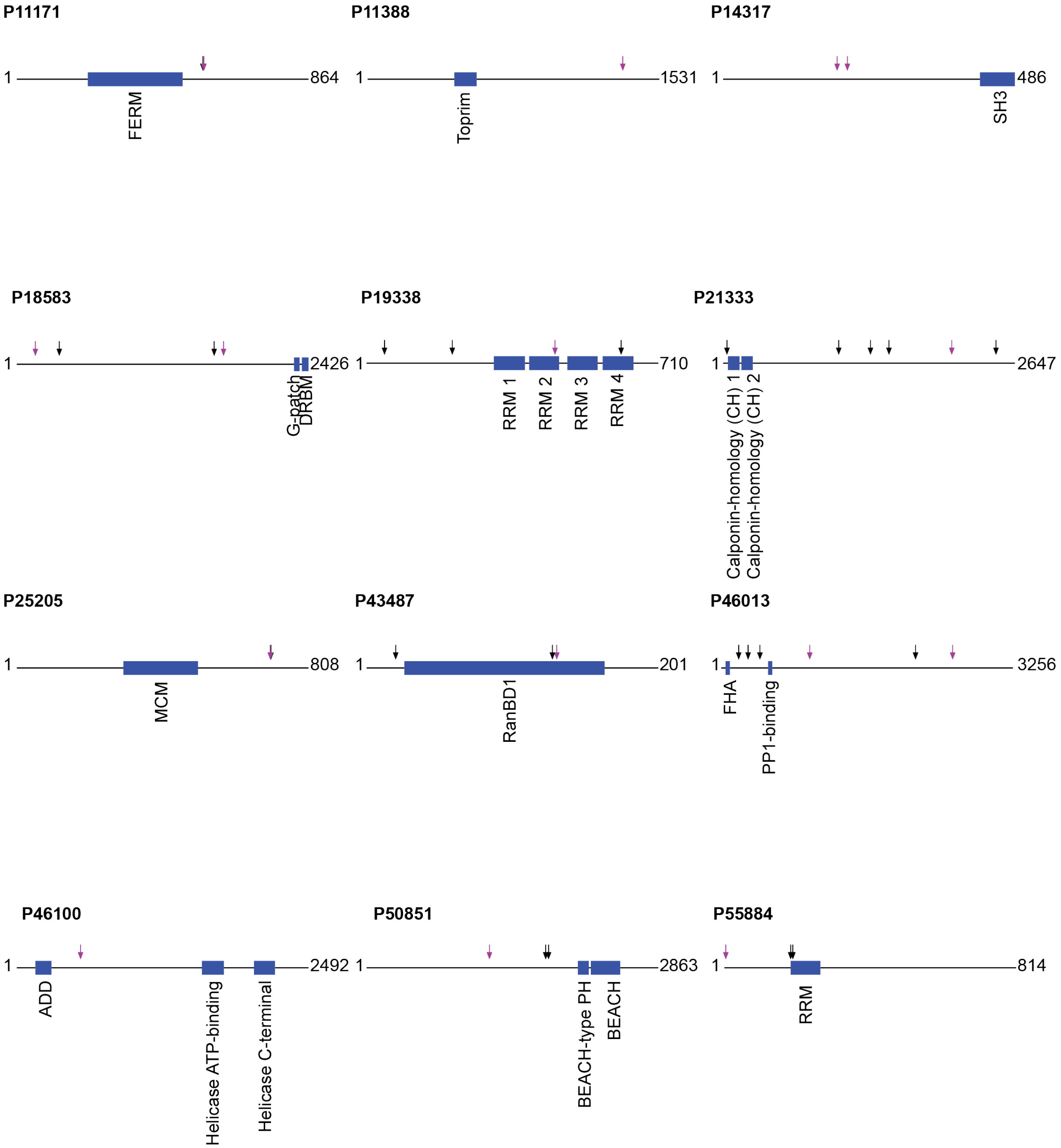

**Figure S22 (cont'd).**

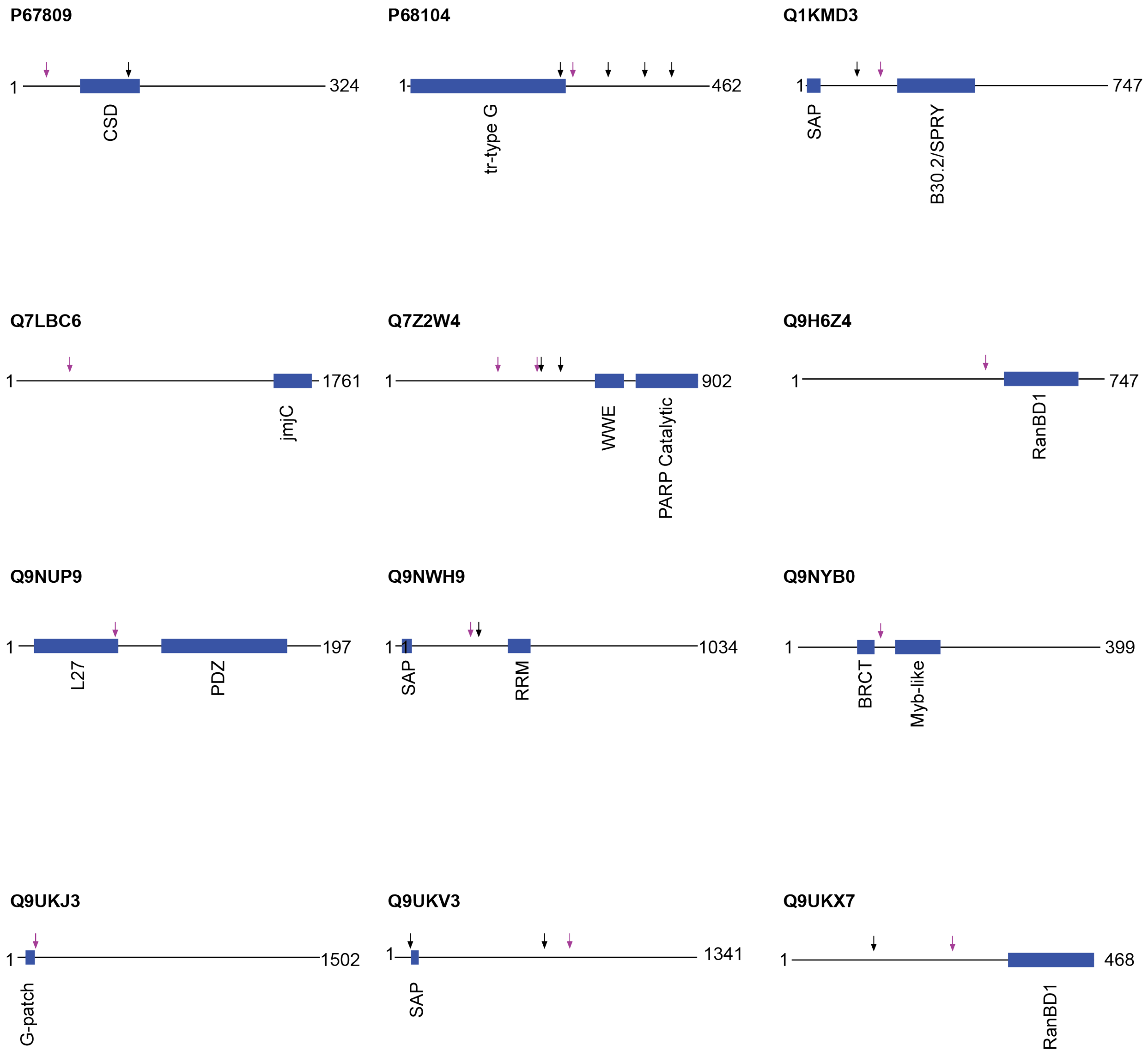

**Figure S22 (cont'd).**

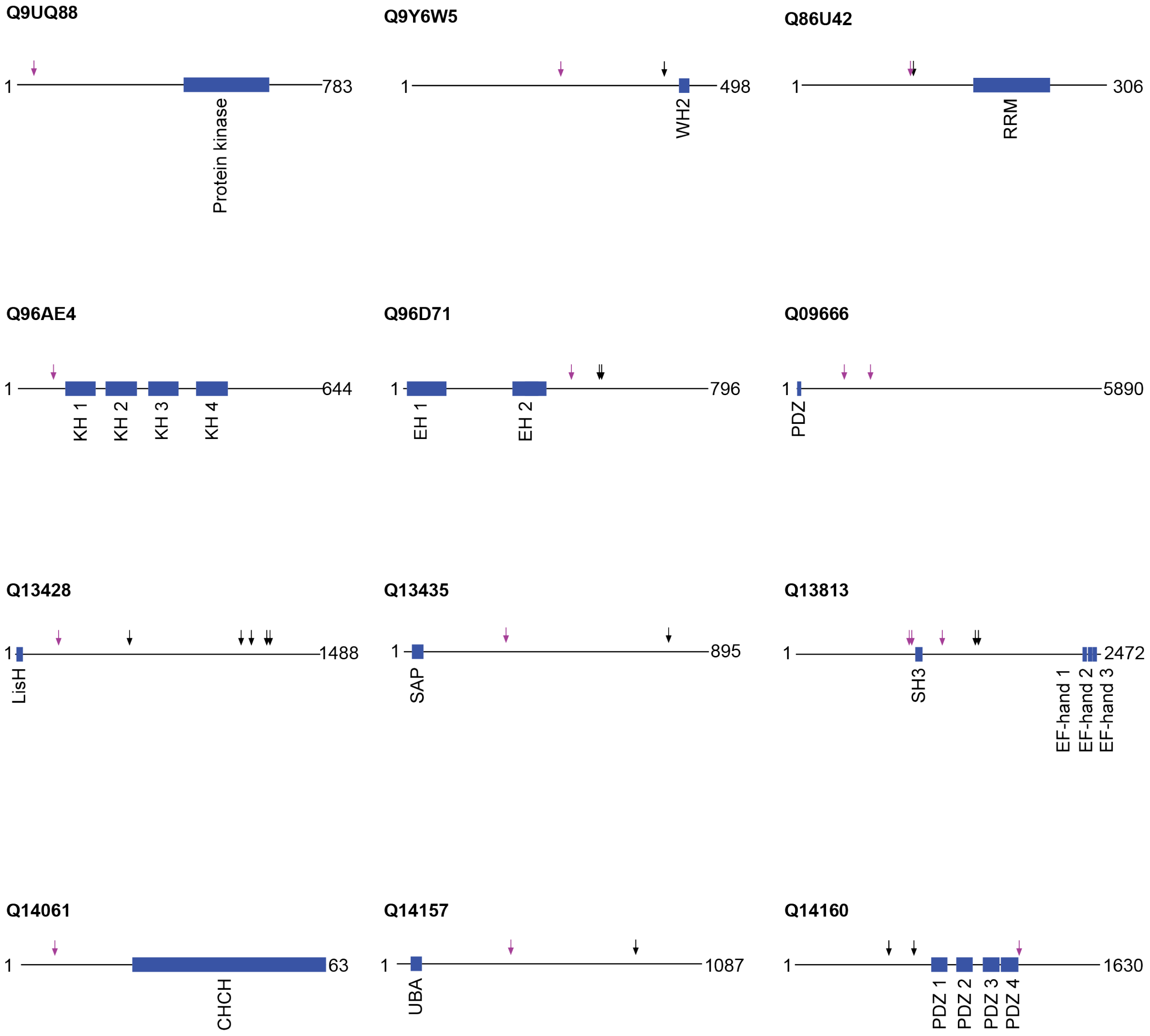

**Figure S22 (cont'd).**

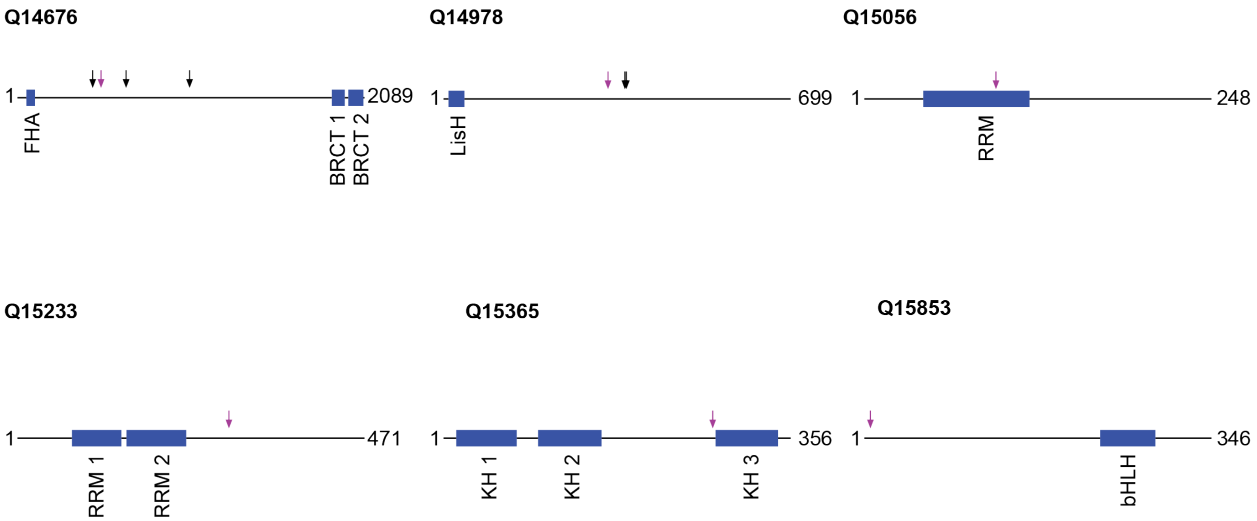

**Figure S23. Frequency of amino acids observed among P1 = D N termini in etoposide-treated cells captured by subtiligase and CHOPPER.** (A) Comparison between subtiligase and CHOPPER. (B) Comparison between a cocktail of subtiligase specificity mutants and CHOPPER.

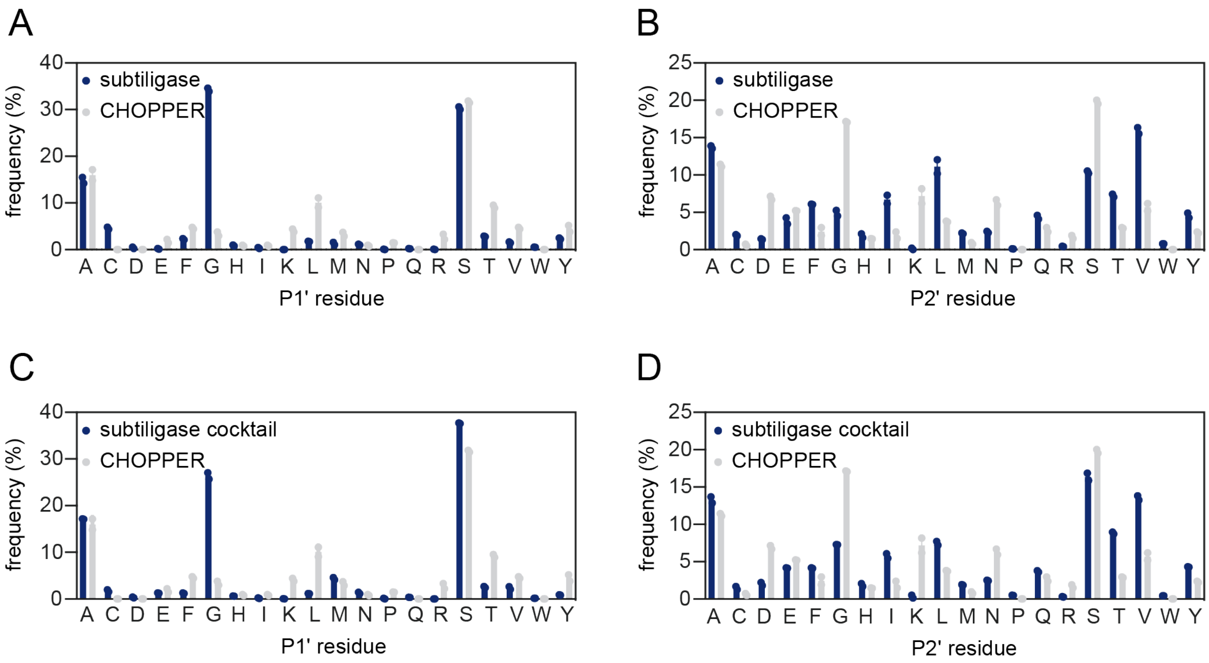

**Figure S24. Structures of chemical probes used in this study.**

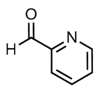

**1**
2-pyridinecarboxaldehyde (2PCA)

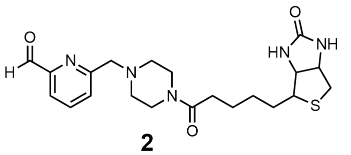

**2**
biotin-2PCA

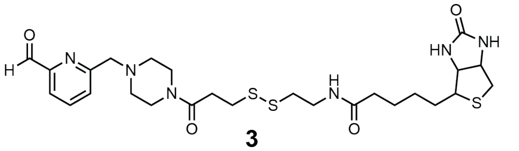

**3**
biotin-SS-2PCA

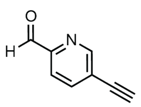

**4**
alkyne-2PCA

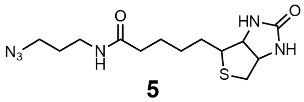

**5**
biotin azide

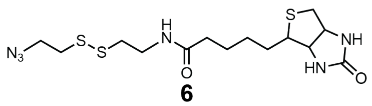

**6**
biotin-disulfide-azide

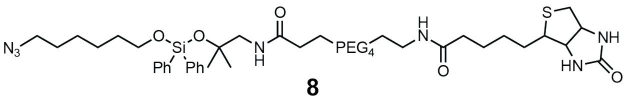

**8**
biotin-DADPS-azide

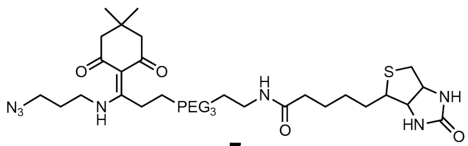

**7**
biotin-Dde-azide

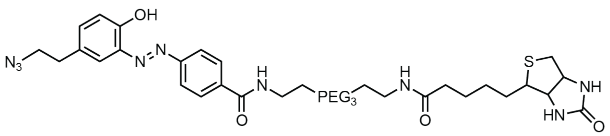

**9**
biotin-Diazo-azide

**Figure S25. Negative mycoplasma contamination tests for HEK293T and Jurkat cells used in this study.** Left, HEK293T; right, Jurkat.

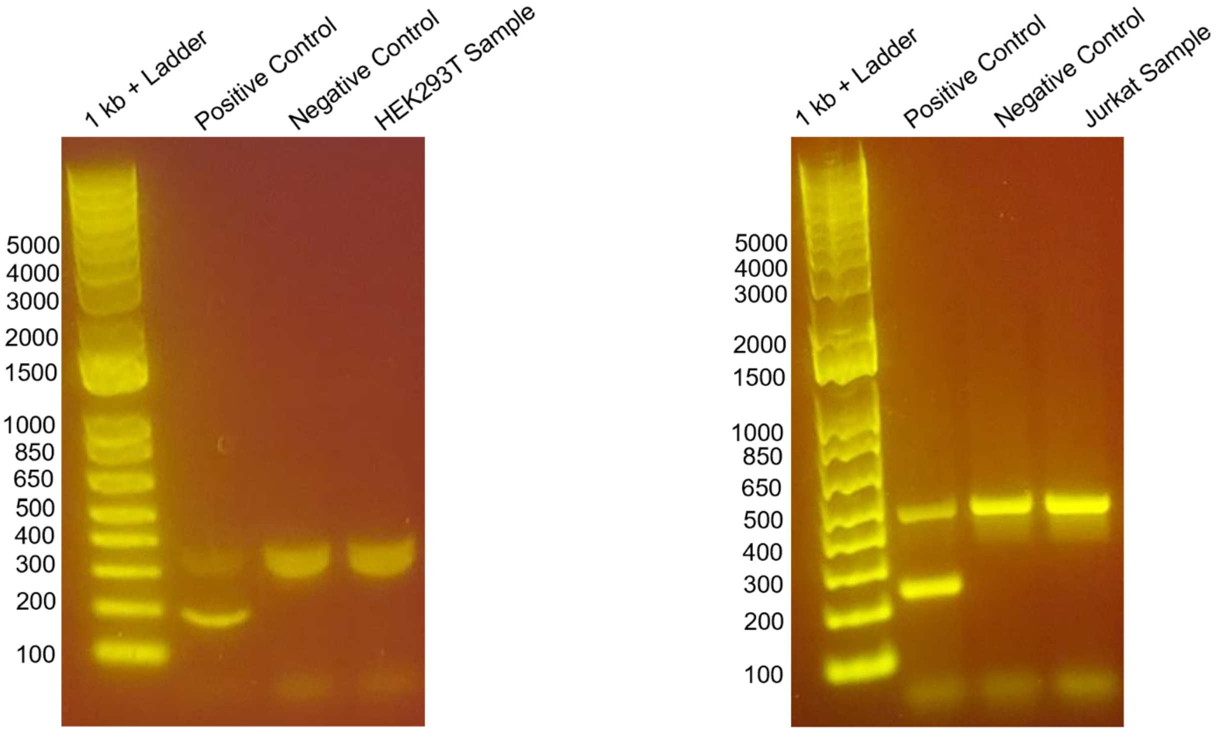

**Figure S26. Structures of 2PCA reagent peptide modifications.** An Ala-Ala peptide is shown as an example.

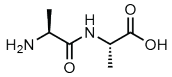

unmodified peptide

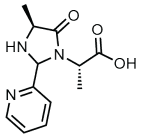

2PCA modification

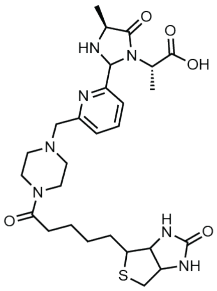

biotin-2PCA modification

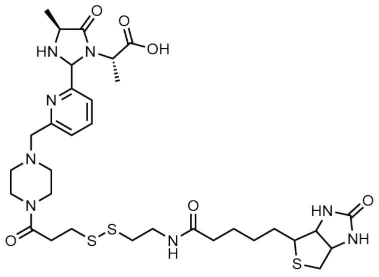

biotin-SS-2PCA modification

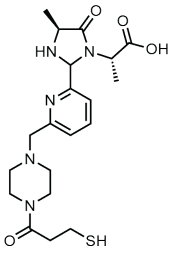

cleaved biotin-SS-2PCA modification

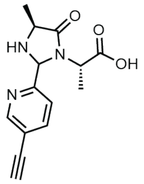

alkyne-2PCA modification

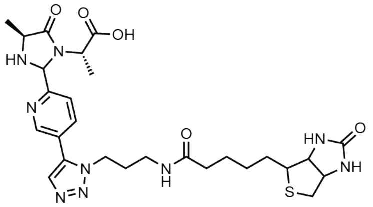

alkyne-2PCA + biotin azide modification

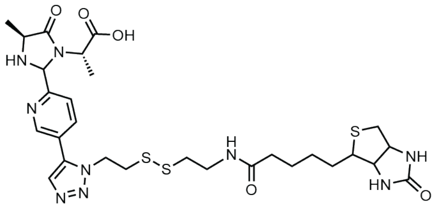

alkyne-2PCA + biotin-SS-azide modification

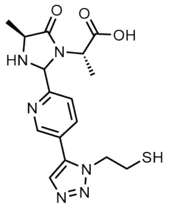

cleaved alkyne-2PCA + biotin-SS-azide modification

### Figure S26 (cont'd).

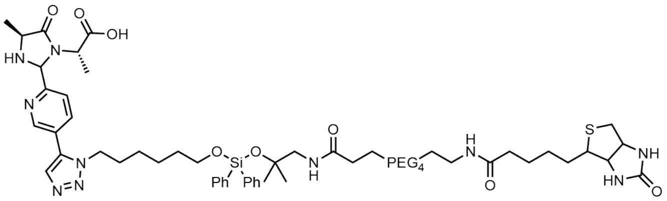

alkyne-2PCA + biotin-DADPS-azide modification

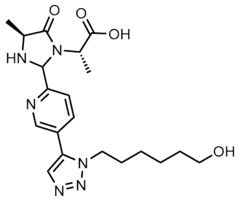

cleaved alkyne-2PCA + biotin-DADPS-azide modification

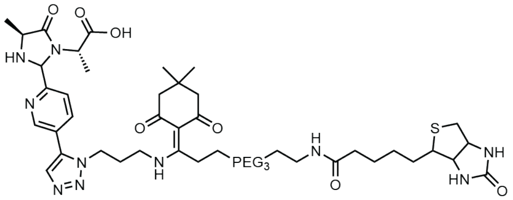

alkyne-2PCA + biotin-Dde-azide modification

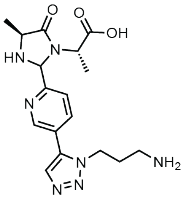

cleaved alkyne-2PCA + biotin-Dde-azide modification

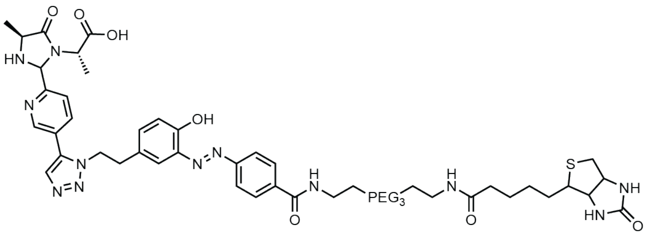

alkyne-2PCA + biotin-Diazo-azide modification

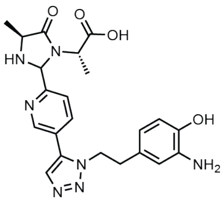

cleaved alkyne-2PCA + biotin-Diazo-azide modification

**Table S1.** Newly identified P1 = D cleavage sites and proteins in etoposide-treated Jurkat cells.

| Uniprot ID | P1' position | protein observed previously? | site observed previously? |
|---|---|---|---|
| Q8NFC6 | 2327 | yes | no |
| Q14160 | 1198 | yes | no |
| O75717 | 865 | no | no |
| P55884 | 4 | yes | no |
| Q16643 | 126 | yes | no |
| Q6IS14 | 12 | no | no |
| O00273 | 7 | yes | no |
| P21333 | 2075 | yes | no |
| Q96AE4 | 75 | no | no |
| Q9H6Z4 | 344 | no | no |
| Q13435 | 300 | yes | no |
| Q86WB0 | 296 | yes | no |
| Q9Y6W5 | 243 | yes | no |
| Q15365 | 276 | no | no |
| P49006 | 56 | yes | no |
| P43487 | 131 | yes | no |
| O14497 | 1786 | yes | no |
| P11388 | 1339 | no | no |
| Q9NWH9 | 257 | yes | no |
| Q6IS14 | 7 | no | no |
| P46013 | 2562 | yes | no |
| Q99848 | 212 | no | no |
| P07910 | 128 | yes | no |
| P25205 | 702 | yes | no |
| P07602 | 406 | yes | no |
| P63267 | 246 | no | no |
| P19338 | 456 | yes | no |
| P46100 | 544 | no | no |
| Q7Z2W4 | 422 | yes | no |
| O75410 | 162 | no | no |
| Q9NUP9 | 63 | no | no |
| P49750 | 937 | no | no |
| Q969S3 | 125 | no | no |
| P52566 | 40 | no | no |
| Q9UKX7 | 249 | yes | no |
| P78559 | 837 | yes | no |
| P14317 | 207 | no | no |
| Q14676 | 503 | yes | no |
| P08575 | 1254 | no | no |
| O60841 | 19 | yes | no |
| Q6UN15 | 157 | yes | no |
| P07910 | 124 | yes | no |
| P07437 | 68 | yes | no |
| Q9H7N4 | 463 | no | no |
| P50851 | 1205 | yes | no |
| Q9BXP5 | 142 | yes | no |
| Q14157 | 406 | yes | no |
| P46013 | 967 | yes | no |
| Q9C0B0 | 524 | no | no |
| Q13573 | 132 | no | no |
| Q13813 | 939 | yes | no |
| Q5H9R7 | 677 | yes | no |
| O95218 | 180 | yes | no |
| P27816 | 152 | yes | no |
| P08195 | 110 | no | no |
| P07602 | 194 | yes | no |

| | | | |
|---|---|---|---|
| P27797 | 211 | yes | no |
| Q86U42 | 109 | yes | no |
| Q15853 | 6 | no | no |
| Q7Z2W4 | 305 | yes | no |
| P48634 | 432 | no | no |
| P47914 | 95 | no | no |
| P07910 | 135 | yes | no |
| Q15233 | 287 | no | no |
| Q3YEC7 | 401 | yes | no |
| P11171 | 554 | yes | no |
| Q96F63 | 53 | no | no |
| Q1KMD3 | 185 | yes | no |
| P28715 | 305 | yes | no |
| P18583 | 154 | yes | no |
| P18583 | 1719 | yes | no |
| Q9Y4E1 | 697 | no | no |
| Q9Y2D5 | 473 | yes | no |
| P49790 | 940 | no | no |
| O75995 | 57 | yes | no |
| Q9NY27 | 199 | no | no |
| P52566 | 20 | no | no |
| Q9UQ88 | 43 | no | no |
| Q13813 | 1186 | yes | no |
| Q14686 | 1876 | no | no |
| Q15056 | 94 | no | no |
| P08670 | 430 | yes | no |
| Q08945 | 497 | no | no |
| Q7LBC6 | 311 | no | no |
| Q96D71 | 439 | yes | no |
| Q14061 | 7 | no | no |
| O00541 | 371 | yes | no |
| Q9UQ35 | 2259 | yes | no |
| Q8IXQ4 | 140 | no | no |
| Q8NFC6 | 1280 | yes | no |
| O15027 | 102 | yes | no |
| Q14978 | 331 | yes | no |
| Q16643 | 457 | yes | no |
| Q8NFC6 | 2897 | yes | no |
| P60866 | 6 | no | no |
| Q02446 | 344 | no | no |
| Q9NYB0 | 109 | no | no |
| P67809 | 25 | yes | no |
| Q9UQ35 | 977 | yes | no |
| Q9UKV3 | 776 | yes | no |
| Q6UN15 | 447 | yes | no |
| Q13428 | 212 | yes | no |
| P53990 | 198 | no | no |
| Q09666 | 1425 | no | no |
| Q09666 | 920 | no | no |
| P68104 | 253 | yes | no |
| Q9UKJ3 | 90 | no | no |
| P14317 | 190 | no | no |
| O00148 | 13 | no | no |
| Q13813 | 918 | yes | no |
| Q9Y520 | 328 | yes | no |
| P04406 | 240 | no | no |

**Table S2.** Monoisotopic masses for peptide modifications used in database searches.

| modification | modification mass | name in supp. dataset |
|---|---|---|
| 2PCA | 89.0265 | 2PCA |
| alkyne-2PCA | 113.0265 | alkyne-2PCA |
| biotin-2PCA | 413.1885 | biotin-pip-2PCA |
| cleaved biotin-disulfide-2PCA | 275.1092 | reduced biotinSS2PCA |
| alkyne-2PCA clicked to biotin azide | 439.1790 | clicked biotin 2PCA |
| cleaved alkyne-2PCA clicked to biotin-SS-azide | 216.0470 | clicked DTT 2PCA |
| cleaved alkyne-2PCA clicked to biotin-DADPS-azide | 256.1324 | Cleaved DADPS biotin azide 2PCA |
| cleaved alkyne-2PCA clicked to biotin-Dde-azide | 213.1014 | Cleaved Dde biotin azide 2PCA |
| cleaved alkyne-2PCA clicked to biotin-Diazo-azide | 291.1120 | Cleaved diazo biotin azide |
| Thioacylation | 87.9983 | Thioacylation |

**Table S3.** List of ProteomeXchange accession numbers and experimental information.

| raw file name | experiment description | source data for | supplementary dataset # | ProteomeXchange # |
|---|---|---|---|---|
| E20220309-02 | E. coli proteome-derived peptide library generated with chymotrypsin | Fig. S1 | Dataset S1 | PXD040044 |
| E20220309-04 | E. coli proteome-derived peptide library generated with GluC | Fig. S1 | Dataset S1 | PXD040044 |
| E20220309-06 | E. coli proteome-derived peptide library generated with trypsin | Fig. S1 | Dataset S1 | PXD040044 |
| E20210730-15 | E. coli proteome-derived peptide library generated with trypsin; treated with 10 mM 2PCA for 4 h at 37C | Fig. 1CDEFG; Fig. S2; Fig. S4 | Dataset S2 | PXD040045 |
| E20210828-15 | E. coli proteome-derived peptide library generated with GluC; treated with 10 mM 2PCA for 4 h at 37C | Fig. 1CDEFG; Fig. S2; Fig. S4 | Dataset S2 | PXD040045 |
| E20210902-21 | E. coli proteome-derived peptide library generated with chymotrypsin; treated with 10 mM 2PCA for 4 h at 37C | Fig. 1CDEFG; Fig. S2; Fig. S4 | Dataset S2 | PXD040045 |
| E20211223-03 | 50 mM 2PCA reacted at 37C for 4 hours in 50 mM phosphate buffer with E. coli chymotrypsin library | Fig. 2B | Dataset S3 | PXD040046 |
| E20211223-05 | 50 mM 2PCA reacted at 37C for 4 hours in 50 mM phosphate buffer with E. coli GluC library | Fig. 2B | Dataset S3 | PXD040046 |
| E20211223-07 | 50 mM 2PCA reacted at 37C for 4 hours in 50 mM phosphate buffer with E. coli trypsin library | Fig. 2B | Dataset S3 | PXD040046 |
| E20211223-09 | 50 mM 2PCA reacted at 45C for 4 hours in 50 mM phosphate buffer with E. coli chymotrypsin library | Fig. 2B | Dataset S3 | PXD040046 |
| E20211223-11 | 50 mM 2PCA reacted at 45C for 4 hours in 50 mM phosphate buffer with E. coli GluC library | Fig. 2B | Dataset S3 | PXD040046 |
| E20211223-13 | 50 mM 2PCA reacted at 45C for 4 hours in 50 mM phosphate buffer with E. coli trypsin library | Fig. 2B | Dataset S3 | PXD040046 |
| E20211223-15 | 50 mM 2PCA reacted at 55C for 4 hours in 50 mM phosphate buffer with E. coli chymotrypsin library | Fig. 2B | Dataset S3 | PXD040046 |
| E20211223-17 | 50 mM 2PCA reacted at 55C for 4 hours in 50 mM phosphate buffer with E. coli GluC library | Fig. 2B | Dataset S3 | PXD040046 |
| E20211223-19 | 50 mM 2PCA reacted at 55C for 4 hours in 50 mM phosphate buffer with E. coli trypsin library | Fig. 2B | Dataset S3 | PXD040046 |

| | | | | |
|---|---|---|---|---|
| E20211227-03 | 50 mM 2PCA reacted at 65C for 4 hours in 50 mM phosphate buffer with E. coli chymotrypsin library | Fig. 2B | Dataset S3 | PXD040046 |
| E20211227-05 | 50 mM 2PCA reacted at 65C for 4 hours in 50 mM phosphate buffer with E. coli GluC library | Fig. 2B | Dataset S3 | PXD040046 |
| E20211227-07 | 50 mM 2PCA reacted at 65C for 4 hours in 50 mM phosphate buffer with E. coli trypsin library | Fig. 2B | Dataset S3 | PXD040046 |
| E20211227-09 | 50 mM 2PCA reacted at 75C for 4 hours in 50 mM phosphate buffer with E. coli chymotrypsin library | Fig. 2B | Dataset S3 | PXD040046 |
| E20211227-11 | 50 mM 2PCA reacted at 75C for 4 hours in 50 mM phosphate buffer with E. coli GluC library | Fig. 2B | Dataset S3 | PXD040046 |
| E20211227-13 | 50 mM 2PCA reacted at 75C for 4 hours in 50 mM phosphate buffer with E. coli trypsin library | Fig. 2B | Dataset S3 | PXD040046 |
| E20211227-30 | 50 mM 2PCA reacted at 37C for 4 hours in 10 mM phosphate buffer with E. coli chymotrypsin library | Fig. 2B | Dataset S3 | PXD040046 |
| E20211227-32 | 50 mM 2PCA reacted at 37C for 4 hours in 10 mM phosphate buffer with E. coli GluC library | Fig. 2B | Dataset S3 | PXD040046 |
| E20211227-34 | 50 mM 2PCA reacted at 37C for 4 hours in 10 mM phosphate buffer with E. coli trypsin library | Fig. 2B | Dataset S3 | PXD040046 |
| E20220114-18 | 50 mM 2PCA reacted at 45C for 4 hours in 10 mM phosphate buffer with E. coli chymotrypsin library | Fig. 2B | Dataset S3 | PXD040046 |
| E20220114-20 | 50 mM 2PCA reacted at 45C for 4 hours in 10 mM phosphate buffer with E. coli GluC library | Fig. 2B | Dataset S3 | PXD040046 |
| E20220114-22 | 50 mM 2PCA reacted at 45C for 4 hours in 10 mM phosphate buffer with E. coli trypsin library | Fig. 2B | Dataset S3 | PXD040046 |
| E20220129-08 | 50 mM 2PCA reacted at 55C for 4 hours in 10 mM phosphate buffer with E. coli chymotrypsin library | Fig. 2B | Dataset S3 | PXD040046 |
| E20220129-10 | 50 mM 2PCA reacted at 55C for 4 hours in 10 mM phosphate buffer with E. coli GluC library | Fig. 2B | Dataset S3 | PXD040046 |
| E20220129-12 | 50 mM 2PCA reacted at 55C for 4 hours in 10 mM phosphate buffer with E. coli trypsin library | Fig. 2B | Dataset S3 | PXD040046 |

| | | | | |
|---|---|---|---|---|
| E20220211-05 | 50 mM 2PCA reacted at 65C for 4 hours in 10 mM phosphate buffer with E. coli GluC library | Fig. 2B | Dataset S3 | PXD040046 |
| E20220211-07 | 50 mM 2PCA reacted at 65C for 4 hours in 10 mM phosphate buffer with E. coli trypsin library | Fig. 2B | Dataset S3 | PXD040046 |
| E20220211-09 | 50 mM 2PCA reacted at 75C for 4 hours in 10 mM phosphate buffer with E. coli chymotrypsin library | Fig. 2B | Dataset S3 | PXD040046 |
| E20220211-11 | 50 mM 2PCA reacted at 75C for 4 hours in 10 mM phosphate buffer with E. coli GluC library | Fig. 2B | Dataset S3 | PXD040046 |
| E20220211-13 | 50 mM 2PCA reacted at 75C for 4 hours in 10 mM phosphate buffer with E. coli trypsin library | Fig. 2B | Dataset S3 | PXD040046 |
| E20220302-10 | 50 mM 2PCA reacted at 65C for 4 hours in 10 mM phosphate buffer with E. coli chymotrypsin library | Fig. 2B | Dataset S3 | PXD040046 |
| E20211227-44 | 50 mM 2PCA reacted at 37C for 20 hours in 50 mM phosphate buffer with E. coli chymotrypsin library | Fig. 2C | Dataset S4 | PXD040047 |
| E20211227-46 | 50 mM 2PCA reacted at 37C for 20 hours in 50 mM phosphate buffer with E. coli GluC library | Fig. 2C | Dataset S4 | PXD040047 |
| E20211227-48 | 50 mM 2PCA reacted at 37C for 20 hours in 50 mM phosphate buffer with E. coli trypsin library | Fig. 2C | Dataset S4 | PXD040047 |
| E20220129-02 | 50 mM 2PCA reacted at 37C for 20 hours in 10 mM phosphate buffer with E. coli chymotrypsin library | Fig. 2C | Dataset S4 | PXD040047 |
| E20220129-04 | 50 mM 2PCA reacted at 37C for 20 hours in 10 mM phosphate buffer with E. coli GluC library | Fig. 2C | Dataset S4 | PXD040047 |
| E20220129-06 | 50 mM 2PCA reacted at 37C for 20 hours in 10 mM phosphate buffer with E. coli trypsin library | Fig. 2C | Dataset S4 | PXD040047 |
| E20220129-14 | 50 mM 2PCA reacted at 75C for 20 hours in 50 mM phosphate buffer with E. coli chymotrypsin library | Fig. 2C | Dataset S4 | PXD040047 |
| E20220129-16 | 50 mM 2PCA reacted at 75C for 20 hours in 50 mM phosphate buffer with E. coli GluC library | Fig. 2C | Dataset S4 | PXD040047 |
| E20220129-18 | 50 mM 2PCA reacted at 75C for 20 hours in 50 mM phosphate buffer with E. coli trypsin library | Fig. 2C | Dataset S4 | PXD040047 |

| | | | | |
|---|---|---|---|---|
| E20220211-15 | 50 mM 2PCA reacted at 65C for 20 hours in 10 mM phosphate buffer with E. coli chymotrypsin library | Fig. 2C | Dataset S4 | PXD040047 |
| E20220211-17 | 50 mM 2PCA reacted at 65C for 20 hours in 10 mM phosphate buffer with E. coli GluC library | Fig. 2C | Dataset S4 | PXD040047 |
| E20220211-19 | 50 mM 2PCA reacted at 65C for 20 hours in 10 mM phosphate buffer with E. coli trypsin library | Fig. 2C | Dataset S4 | PXD040047 |
| E20220211-21 | 50 mM 2PCA reacted at 75C for 20 hours in 10 mM phosphate buffer with E. coli chymotrypsin library | Fig. 2C | Dataset S4 | PXD040047 |
| E20220211-23 | 50 mM 2PCA reacted at 75C for 20 hours in 10 mM phosphate buffer with E. coli GluC library | Fig. 2C | Dataset S4 | PXD040047 |
| E20220218-10 | 50 mM 2PCA reacted at 45C for 20 hours in 10 mM phosphate buffer with E. coli chymotrypsin library | Fig. 2C | Dataset S4 | PXD040047 |
| E20220218-12 | 50 mM 2PCA reacted at 45C for 20 hours in 10 mM phosphate buffer with E. coli GluC library | Fig. 2C | Dataset S4 | PXD040047 |
| E20220218-14 | 50 mM 2PCA reacted at 45C for 20 hours in 10 mM phosphate buffer with E. coli trypsin library | Fig. 2C | Dataset S4 | PXD040047 |
| E20220218-16 | 50 mM 2PCA reacted at 55C for 20 hours in 10 mM phosphate buffer with E. coli chymotrypsin library | Fig. 2C | Dataset S4 | PXD040047 |
| E20220218-18 | 50 mM 2PCA reacted at 55C for 20 hours in 10 mM phosphate buffer with E. coli GluC library | Fig. 2C | Dataset S4 | PXD040047 |
| E20220218-20 | 50 mM 2PCA reacted at 55C for 20 hours in 10 mM phosphate buffer with E. coli trypsin library | Fig. 2C | Dataset S4 | PXD040047 |
| E20220225-03 | 50 mM 2PCA reacted at 45C for 20 hours in 50 mM phosphate buffer with E. coli chymotrypsin library | Fig. 2C | Dataset S4 | PXD040047 |
| E20220225-05 | 50 mM 2PCA reacted at 45C for 20 hours in 50 mM phosphate buffer with E. coli GluC library | Fig. 2C | Dataset S4 | PXD040047 |
| E20220225-07 | 50 mM 2PCA reacted at 45C for 20 hours in 50 mM phosphate buffer with E. coli trypsin library | Fig. 2C | Dataset S4 | PXD040047 |
| E20220225-09 | 50 mM 2PCA reacted at 55C for 20 hours in 50 mM phosphate buffer with E. coli chymotrypsin library | Fig. 2C | Dataset S4 | PXD040047 |

| | | | | |
|---|---|---|---|---|
| E20220225-11 | 50 mM 2PCA reacted at 55C for 20 hours in 50 mM phosphate buffer with E. coli GluC library | Fig. 2C | Dataset S4 | PXD040047 |
| E20220225-13 | 50 mM 2PCA reacted at 55C for 20 hours in 50 mM phosphate buffer with E. coli trypsin library | Fig. 2C | Dataset S4 | PXD040047 |
| E20220302-04 | 50 mM 2PCA reacted at 65C for 20 hours in 50 mM phosphate buffer with E. coli chymotrypsin library | Fig. 2C | Dataset S4 | PXD040047 |
| E20220302-06 | 50 mM 2PCA reacted at 65C for 20 hours in 50 mM phosphate buffer with E. coli GluC library | Fig. 2C | Dataset S4 | PXD040047 |
| E20220302-08 | 50 mM 2PCA reacted at 65C for 20 hours in 50 mM phosphate buffer with E. coli trypsin library | Fig. 2C | Dataset S4 | PXD040047 |
| E20220302-12 | 50 mM 2PCA reacted at 75C for 20 hours in 10 mM phosphate buffer with E. coli trypsin library | Fig. 2C | Dataset S4 | PXD040047 |
| E20201217-29 | 30 mM 2PCA reacted at 37C for 20 hours in 10 mM phosphate buffer with E. coli chymotrypsin library | Fig. 2DE, Fig. S5, Fig. S6 | Dataset S5 | PXD040053 |
| E20201217-35 | 15 mM 2PCA reacted at 37C for 20 hours in 10 mM phosphate buffer with E. coli chymotrypsin library | Fig. 2DE, Fig. S5, Fig. S6 | Dataset S5 | PXD040053 |
| E20201217-39 | 15 mM 2PCA reacted at 37C for 20 hours in 10 mM phosphate buffer with E. coli trypsin library | Fig. 2DE, Fig. S5, Fig. S6 | Dataset S5 | PXD040053 |
| E20201217-41 | 10 mM 2PCA reacted at 37C for 20 hours in 10 mM phosphate buffer with E. coli chymotrypsin library | Fig. 2DE, Fig. S5, Fig. S6 | Dataset S5 | PXD040053 |
| E20210104-08 | 5 mM 2PCA reacted at 37C for 20 hours in 10 mM phosphate buffer with E. coli chymotrypsin library | Fig. 2DE, Fig. S5, Fig. S6 | Dataset S5 | PXD040053 |
| E20210112-03 | 50 mM 2PCA reacted at 37C for 20 hours in 10 mM phosphate buffer with E. coli chymotrypsin library | Fig. 2DE, Fig. S5, Fig. S6 | Dataset S5 | PXD040053 |
| E20210112-07 | 50 mM 2PCA reacted at 37C for 20 hours in 10 mM phosphate buffer with E. coli trypsin library | Fig. 2DE, Fig. S5, Fig. S6 | Dataset S5 | PXD040053 |
| E20210113-03 | 10 mM 2PCA reacted at 37C for 20 hours in 10 mM phosphate buffer with E. coli trypsin library | Fig. 2DE, Fig. S5, Fig. S6 | Dataset S5 | PXD040053 |
| E20210113-05 | 5 mM 2PCA reacted at 37C for 20 hours in 10 mM phosphate buffer with E. coli trypsin library | Fig. 2DE, Fig. S5, Fig. S6 | Dataset S5 | PXD040053 |

| | | | | |
|---|---|---|---|---|
| E20210127-03 | 0.01 mM 2PCA reacted at 37C for 20 hours in 10 mM phosphate buffer with E. coli chymotrypsin library | Fig. 2DE, Fig. S5, Fig. S6 | Dataset S5 | PXD040053 |
| E20210127-05 | 0.01 mM 2PCA reacted at 37C for 20 hours in 10 mM phosphate buffer with E. coli trypsin library | Fig. 2DE, Fig. S5, Fig. S6 | Dataset S5 | PXD040053 |
| E20210202-05 | 5 mM 2PCA reacted at 37C for 20 hours in 10 mM phosphate buffer with E. coli GluC library | Fig. 2DE, Fig. S5, Fig. S6 | Dataset S5 | PXD040053 |
| E20210202-07 | 10 mM 2PCA reacted at 37C for 20 hours in 10 mM phosphate buffer with E. coli GluC library | Fig. 2DE, Fig. S5, Fig. S6 | Dataset S5 | PXD040053 |
| E20210202-09 | 15 mM 2PCA reacted at 37C for 20 hours in 10 mM phosphate buffer with E. coli GluC library | Fig. 2DE, Fig. S5, Fig. S6 | Dataset S5 | PXD040053 |
| E20210202-11 | 30 mM 2PCA reacted at 37C for 20 hours in 10 mM phosphate buffer with E. coli GluC library | Fig. 2DE, Fig. S5, Fig. S6 | Dataset S5 | PXD040053 |
| E20210202-13 | 50 mM 2PCA reacted at 37C for 20 hours in 10 mM phosphate buffer with E. coli GluC library | Fig. 2DE, Fig. S5, Fig. S6 | Dataset S5 | PXD040053 |
| E20210507-03 | 0.025 mM 2PCA reacted at 37C for 20 hours in 10 mM phosphate buffer with E. coli chymotrypsin library | Fig. 2DE, Fig. S5, Fig. S6 | Dataset S5 | PXD040053 |
| E20210507-07 | 0.1 mM 2PCA reacted at 37C for 20 hours in 10 mM phosphate buffer with E. coli chymotrypsin library | Fig. 2DE, Fig. S5, Fig. S6 | Dataset S5 | PXD040053 |
| E20210507-09 | 1 mM 2PCA reacted at 37C for 20 hours in 10 mM phosphate buffer with E. coli chymotrypsin library | Fig. 2DE, Fig. S5, Fig. S6 | Dataset S5 | PXD040053 |
| E20210507-11 | 2 mM 2PCA reacted at 37C for 20 hours in 10 mM phosphate buffer with E. coli chymotrypsin library | Fig. 2DE, Fig. S5, Fig. S6 | Dataset S5 | PXD040053 |
| E20210507-13 | 0.025 mM 2PCA reacted at 37C for 20 hours in 10 mM phosphate buffer with E. coli GluC library | Fig. 2DE, Fig. S5, Fig. S6 | Dataset S5 | PXD040053 |
| E20210507-15 | 0.01 mM 2PCA reacted at 37C for 20 hours in 10 mM phosphate buffer with E. coli GluC library | Fig. 2DE, Fig. S5, Fig. S6 | Dataset S5 | PXD040053 |
| E20210507-17 | 0.1 mM 2PCA reacted at 37C for 20 hours in 10 mM phosphate buffer with E. coli GluC library | Fig. 2DE, Fig. S5, Fig. S6 | Dataset S5 | PXD040053 |
| E20210507-19 | 1 mM 2PCA reacted at 37C for 20 hours in 10 mM phosphate buffer with E. coli GluC library | Fig. 2DE, Fig. S5, Fig. S6 | Dataset S5 | PXD040053 |

| | | | | |
|---|---|---|---|---|
| E20210507-21 | 2 mM 2PCA reacted at 37C for 20 hours in 10 mM phosphate buffer with E. coli GluC library | Fig. 2DE, Fig. S5, Fig. S6 | Dataset S5 | PXD040053 |
| E20210507-23 | 0.025 mM 2PCA reacted at 37C for 20 hours in 10 mM phosphate buffer with E. coli trypsin library | Fig. 2DE, Fig. S5, Fig. S6 | Dataset S5 | PXD040053 |
| E20210507-27 | 0.1 mM 2PCA reacted at 37C for 20 hours in 10 mM phosphate buffer with E. coli trypsin library | Fig. 2DE, Fig. S5, Fig. S6 | Dataset S5 | PXD040053 |
| E20210927-11 | 30 mM 2PCA reacted at 37C for 20 hours in 10 mM phosphate buffer with E. coli trypsin library | Fig. 2DE, Fig. S5, Fig. S6 | Dataset S5 | PXD040053 |
| E20220817-02 | 1 mM 2PCA reacted at 37C for 20 hours in 10 mM phosphate buffer with E. coli trypsin library | Fig. 2DE, Fig. S5, Fig. S6 | Dataset S5 | PXD040053 |
| E20220817-04 | 2 mM 2PCA reacted at 37C for 20 hours in 10 mM phosphate buffer with E. coli trypsin library | Fig. 2DE, Fig. S5, Fig. S6 | Dataset S5 | PXD040053 |
| E20211109-05 | Human proteome-derived peptide library generated with trypsin; treated with 10 mM alkyne-2PCA for 20 h at 37C | Fig. S7 | Dataset S6 | PXD040048 |
| E20211117-03 | Human proteome-derived peptide library generated with trypsin; treated with 10 mM alkyne-2PCA for 20 h at 37C; clicked with biotin azide | Fig. S8 | Dataset S7 | PXD040049 |
| E20220216-01 | Biotin-SS-2PCA E. coli trypsin library | Fig. 3B | Dataset S8 | PXD040050 |
| E20220301-13 | Biotin-SS-2PCA E. coli GluC library | Fig. 3B | Dataset S8 | PXD040050 |
| E20220301-15 | Biotin-2PCA E. coli trypsin library | Fig. 3B | Dataset S9 | PXD040051 |
| E20220301-17 | Biotin-2PCA E. coli GluC library | Fig. 3B | Dataset S9 | PXD040051 |
| E20220221-14 | Alkyne-2PCA, biotin azide, E. coli trypsin library | Fig. 3C | Dataset S10 | PXD040052 |
| E20220407-03 | Alkyne-2PCA, biotin azide, E. coli GluC library | Fig. 3C | Dataset S10 | PXD040052 |
| E20220221-08 | Alkyne-2PCA, biotin disulfide azide, E. coli trypsin library | Fig. 3C | Dataset S11 | PXD040054 |
| E20220304-02 | Alkyne-2PCA, biotin disulfide azide, E. coli GluC library | Fig. 3C | Dataset S11 | PXD040054 |
| E20220221-12 | Alkyne-2PCA, biotin DADPS azide, E. coli trypsin library | Fig. 3C | Dataset S12 | PXD040055 |
| E20220304-08 | Alkyne-2PCA, biotin DADPS azide, E. coli GluC library | Fig. 3C | Dataset S12 | PXD040055 |
| E20220221-10 | Alkyne-2PCA, biotin Dde azide, E. coli trypsin library | Fig. 3C | Dataset S13 | PXD040056 |
| E20220304-04 | Alkyne-2PCA, biotin Dde azide, E. coli GluC library | Fig. 3C | Dataset S13 | PXD040056 |
| E20220221-06 | Alkyne-2PCA, biotin Diazo azide, E. coli trypsin library | Fig. 3C | Dataset S14 | PXD040057 |
| E20220304-06 | Alkyne-2PCA, biotin Diazo azide, E. coli GluC library | Fig. 3C | Dataset S14 | PXD040057 |

| | | | | |
|---|---|---|---|---|
| E20210711-03 | Human (HEK293T) proteome-derived peptide library generated with trypsin | Fig. S9 | Dataset S15 | PXD040058 |
| E20210711-05 | Human (HEK293T) proteome-derived peptide library generated with chymotrypsin | Fig. S9 | Dataset S15 | PXD040058 |
| E20210711-07 | Human (HEK293T) proteome-derived peptide library generated with GluC | Fig. S9 | Dataset S15 | PXD040058 |
| E20220202-06 | Human (HEK293T) proteome-derived peptide library generated with trypsin; modified with 50 mM 2PCA at 37C overnight | Fig. S10 | Dataset S16 | PXD040059 |
| E20220202-08 | Human (HEK293T) proteome-derived peptide library generated with chymotrypsin; modified with 50 mM 2PCA at 37C overnight | Fig. S10 | Dataset S16 | PXD040059 |
| E20220202-10 | Human (HEK293T) proteome-derived peptide library generated with GluC; modified with 50 mM 2PCA at 37C overnight | Fig. S10 | Dataset S16 | PXD040059 |
| E20220310-02 | E. coli proteome-derived peptide library generated with chymotrypsin; blocked with 50 mM 2PCA at 37C overnight | Fig. S10 | Dataset S16 | PXD040059 |
| E20220310-04 | E. coli proteome-derived peptide library generated with GluC; blocked with 50 mM 2PCA at 37C overnight | Fig. S10 | Dataset S16 | PXD040059 |
| E20220310-06 | E. coli proteome-derived peptide library generated with trypsin; blocked with 50 mM 2PCA at 37C overnight | Fig. S10 | Dataset S16 | PXD040059 |
| E20210517-03 | Trypsin PICS with E. coli GluC library | Fig. 4BCD, Fig. S11 | Dataset S17 | PXD040060 |
| E20210517-05 | Trypsin PICS with E. coli trypsin library | Fig. 4BCD, Fig. S11 | Dataset S17 | PXD040060 |
| E20210517-07 | Trypsin PICS with E. coli chymotrypsin library | Fig. 4BCD, Fig. S11 | Dataset S17 | PXD040060 |
| E20220315-02 | Trypsin PICS2 with E. coli GluC library | Fig. 4BCD, Fig. S11 | Dataset S18 | PXD040061 |
| E20220315-04 | Trypsin PICS2 with E. coli chymotrypsin library | Fig. 4BCD, Fig. S11 | Dataset S18 | PXD040061 |
| E20220315-06 | Trypsin PICS2 with E. coli trypsin library | Fig. 4BCD, Fig. S11 | Dataset S18 | PXD040061 |
| E20220322-02 | LysargiNase PICS2 with E. coli GluC library | Fig. 4E, Fig. S12 | Dataset S19 | PXD040062 |
| E20220322-04 | LysargiNase PICS2 with E. coli chymotrypsin library | Fig. 4E, Fig. S12 | Dataset S19 | PXD040062 |
| E20220322-06 | LysargiNase PICS2 with E. coli trypsin library | Fig. 4E, Fig. S12 | Dataset S19 | PXD040062 |
| E20220414-12 | GluC PICS2 with E. coli trypsin library | Fig. S13 | Dataset S20 | PXD040063 |
| E20220408-07 | Chymotrypsin PICS2 with E. coli trypsin library | Fig. S14 | Dataset S21 | PXD040064 |

| | | | | |
|---|---|---|---|---|
| E20221125-24 | N-terminally dimethylated E. coli chymotrypsin library | Fig. S15 | Dataset S22 | PXD040065 |
| E20221125-26 | N-terminally dimethylated E. coli GluC library | Fig. S15 | Dataset S22 | PXD040065 |
| E20221129-02 | Kex2 PICS2 with N-terminally dimethylated E. coli chymotrypsin library | Fig. 4FG, Fig. S16 | Dataset S23 | PXD040066 |
| E20221129-04 | Kex2 PICS2 with N-terminally dimethylated E. coli GluC library | Fig. 4FG, Fig. S16 | Dataset S23 | PXD040066 |
| E20220327-02 | Furin PICS2 with 2PCA-blocked human GluC library | Fig. 4I, Fig. S18 | Dataset S24 | PXD040067 |
| E20220327-04 | Furin PICS2 with 2PCA-blocked human chymotrypsin library | Fig. 4I, Fig. S18 | Dataset S24 | PXD040067 |
| E20221208-03 | PCSK2 PICS2 with N-terminally dimethylated human chymotrypsin library | Fig. 4J, Fig. S19 | Dataset S25 | PXD040068 |
| E20221208-05 | PCSK2 PICS2 with N-terminally dimethylated human GluC library | Fig. 4J, Fig. S19 | Dataset S25 | PXD040068 |
| E20220729-03 | CHOPPER with etoposide-treated Jurkat cells replicate 1 | Fig. 5 | Dataset S26 | PXD040069 |
| E20220729-05 | CHOPPER with etoposide-treated Jurkat cells replicate 2 | Fig. 5 | Dataset S26 | PXD040069 |
| E20220729-07 | CHOPPER with DMSO-treated Jurkat cells replicate 1 | Fig. 5 | Dataset S27 | PXD040070 |
| E20220729-09 | CHOPPER with DMSO-treated Jurkat cells replicate 2 | Fig. 5 | Dataset S27 | PXD040070 |
